# Single molecule studies reveal branched pathways for activator-dependent assembly of RNA polymerase II pre-initiation complexes

**DOI:** 10.1101/2021.06.20.449130

**Authors:** Inwha Baek, Larry J. Friedman, Jeff Gelles, Stephen Buratowski

## Abstract

RNA polymerase II (Pol II) transcription reconstituted from purified factors suggests pre-initiation complexes (PICs) can assemble by sequential incorporation of factors at the TATA box. However, these basal transcription reactions are generally independent of activators and co-activators. To study PIC assembly under more realistic conditions, we used single-molecule microscopy to visualize factor dynamics during activator-dependent reactions in nuclear extracts. Surprisingly, Pol II, TFIIF, and TFIIE can pre-assemble on enhancer-bound activators before loading into PICs, and multiple Pol II complexes can bind simultaneously to create a localized cluster. Unlike TFIIF and TFIIE, TFIIH binding is singular and dependent on the basal promoter. Activator-tethered factors exhibit dwell times on the order of seconds. In contrast, PICs can persist on the order of minutes in the absence of nucleotide triphosphates, although TFIIE remains unexpectedly dynamic even after TFIIH incorporation. Our kinetic measurements lead to a new branched model for activator-dependent PIC assembly.

**HIGHLIGHTS:** Single molecule microscopy reveals unexpected dynamics of RNA Pol II and GTFs

Multiple Pol IIs cluster on UAS/enhancer-bound activators before binding the core promoter

Pol II, TFIIF, and TFIIE, but not TFIIH, can pre-assemble at the UAS/enhancer

Activators increase the rates of Pol II and GTF association with DNA

**eTOC Blurb:** Single-molecule microscopy experiments by Baek et al. show that RNA polymerase II and basal transcription factors TFIIF and TFIIE preassemble on UAS/enhancer-bound activators, poised for loading into initiation complexes with TFIIH at the core promoter. Transcription activators kinetically enhance factor recruitment, creating a localized cluster of polymerases at the UAS/enhancer.

## INTRODUCTION

Eukaryotic RNA polymerase II (Pol II) transcribes messenger RNAs (mRNAs) and some non-coding RNA species. Pol II initiates at specialized genomic regions called promoters, yet does not recognize specific DNA sequences itself (Nikolov and Burley, 1997). Promoter recognition is conferred by the general transcription factors (GTFs) TFIID, TFIIA, TFIIB, TFIIF, TFIIE, and TFIIH, which assemble Pol II into a preinitiation complex (PIC) (reviewed in (Hahn, 2004; Orphanides et al., 1996; Roeder, 1996; Schier and Taatjes, 2020; Thomas and Chiang, 2006)). Gene expression is often regulated at the level of PIC assembly, yet details of this process remain unclear.

A spectrum of models has been proposed. The prevailing PIC formation model proposes sequential assembly of factors on the core promoter, beginning with DNA recognition by TFIID, or even just its TATA-Binding Protein (TBP) subunit. TFIIA and TFIIB join next via direct TBP contacts (Buratowski et al., 1989). Pol II and TFIIF are recruited next, followed by TFIIE and finally TFIIH (He et al., 2013; Luse, 2014). This model of ordered binding is primarily based on electrophoretic mobility shift and footprinting assays (Buratowski et al., 1989; Inostroza et al., 1991; Peterson et al., 1991), which monitor complexes from the perspective of the DNA template. In these early *in vitro* experiments, PICs were assembled from chromatographically separated and purified GTFs (reviewed in (Thomas and Chiang, 2006)), usually in the absence of transcription activators, Mediator, or the TBP-associated factors (TAFs). While the factor contacts predicted by the stepwise model have been confirmed by structural studies (He et al., 2013; He et al., 2016; Plaschka et al., 2016; Schilbach et al., 2017), it remains unclear whether the partial complexes observed *in vitro* represent physiologically relevant intermediates.

In alternate models, some GTFs form subcomplexes before joining the PIC. For example, TFIIF was first purified as the RNA polymerase II-associated proteins RAP30 and RAP74 (Flores et al., 1990; Flores et al., 1988; Sopta et al., 1985). Similarly, direct physical and functional interactions between TFIIE and TFIIH can be observed *in vitro* (Ohkuma et al., 1995). At the most extreme end of the model spectrum, co-immunoprecipitation studies in both yeast and mammals led to proposals that a Pol II “holoenzyme”, containing multiple GTFs, Mediator, and often other co-activators, arrives at the promoter as a pre-assembled complex (Kim et al., 1994; Koleske and Young, 1994; Maldonado et al., 1996; Ossipow et al., 1995; Ranish et al., 1999).

The previous biochemical and structural experiments underlying PIC models are limited in that they can only isolate and characterize stable complexes. GTF dynamics are rarely directly measured, and when they are the results are typically population-averaged. In contrast, single molecule experiments can detect short-lived intermediates, and can reveal alternative assembly pathways that would otherwise be convoluted in ensemble assays. Recent single molecule papers have shown that TFIIB binds the TBP-promoter complex only transiently until Pol II joins the complex (Zhang et al., 2016), and addressed how TFIIH pulls downstream DNA into the Pol II active site to generate an open complex (Fazal et al., 2015; Tomko et al., 2017).

While important mechanistic insights have emerged, one caveat of the single molecule experiments published to date is that they used purified GTFs, where transcription is independent of the activators and co-activators required for gene expression *in vivo*. *In vivo* single-molecule imaging has therefore emerged as an essential complementary approach for studying factor dynamics, but comes with its own limitations (Zhang and Tjian, 2018). To bridge this gap, we developed a system for fluorescently labeling and visualizing single transcription factor molecules within yeast nuclear extract (Rosen et al., 2020). Containing the full repertoire of nuclear factors, these extracts approximate physiological conditions better than reactions reconstituted with a limited set of purified factors. We and others previously showed that nuclear extracts faithfully recapitulate activator-dependent PIC assembly, initiation, and elongation on bead-immobilized templates (Joo et al., 2019; Joo et al., 2017; Kang et al., 1995; Rani et al., 2004; Ranish et al., 1999; Sikorski et al., 2012; Yudkovsky et al., 2000). Here we study late stage PIC assembly using Colocalization Single Molecule Spectroscopy (CoSMoS), a multi-wavelength single-molecule fluorescence method that measures protein binding at individual DNA molecules tethered on microscope slides (Friedman et al., 2006; Hoskins et al., 2011). The relative dynamics of Pol II and the GTFs TFIIF, TFIIE, and TFIIH on DNA were characterized.

Our results argue against a simple sequential assembly of PICs on the TATA box. We show that one or more Pol II molecules first transiently associate with the upstream activating sequence (UAS) tethered by activator, with subsequent transfer to the core promoter/TATA box. TFIIF primarily arrives with Pol II at the UAS, although a substantial fraction of TFIIF associates after Pol II. TFIIE is recruited after Pol II and TFIIF, but its initial binding is also independent of the core promoter DNA. In marked contrast to TFIIF and TFIIE, association of TFIIH with DNA is completely dependent on the core promoter. Therefore, activator-stimulated transcription produces a branched pathway, where multiple partial complexes containing Pol II, TFIIF, and even TFIIE can pre-assemble on the UAS, locally concentrated to facilitate PIC formation at the core promoter.

## RESULTS

### Fluorescence imaging of individual PICs in nuclear extract

For imaging single molecule kinetics of Pol II and GTFs, we constructed a yeast strain in which Rpb1, the largest subunit of Pol II, is genetically fused at the C-terminus to the SNAP_f_ tag (Keppler et al., 2003). Rpb1-SNAP_f_ was labeled with SNAP-Surface 549 (green-excited dye DY549) during nuclear extract preparation (**Fig. S1A**) as previously described (Rosen et al., 2020). For labeling individual GTFs, the factor of interest was genetically fused to *E. coli* dihydrofolate reductase (DHFR) (Calloway et al., 2007). The GTF^DHFR^ was labeled by adding Cy5-TMP, comprising the DHFR inhibitor trimethoprim (TMP) linked to the red-excited dye Cy5, to reactions immediately prior to imaging (Hoskins et al., 2011). The SNAP_f_/DHFR double-fusion strains (**Table S1**) had normal growth and *in vitro* transcription activity relative to untagged parent strains (**Fig. S1C**). The fusions did not appreciably perturb protein expression levels (**Fig. S1D**).

The transcription template, hereafter referred to as UAS+promoter, has a five Gal4-binding site UAS linked to the *CYC1* core promoter (Joo et al., 2019; Joo et al., 2017) (**Fig. 1A**, top). The downstream end of the 299 bp fragment was labeled with the blue-excited dye Alexa Fluor (AF488), while the other end was biotinylated for tethering to the microscope slide surface (**Fig. 1B**). DNAs were visualized by micromirror multi-wavelength total internal reflection fluorescence (TIRF) microscopy (Friedman et al., 2006). Hundreds of DNA molecules can be simultaneously monitored in a given field-of-view (**Fig. 1C**). After marking positions of the UAS+promoter template molecules, a second DNA carrying only the UAS (UAS, **Fig. 1A**, bottom) was similarly tethered to the same slide surface and imaged (Friedman et al., 2013) to serve as an internal negative control for core promoter-specific binding. To account for non-specific background binding to the slide, at least twice as many locations lacking DNA fluorescence (“off DNA”) as DNA locations were randomly selected as controls.

**Figure 1.**
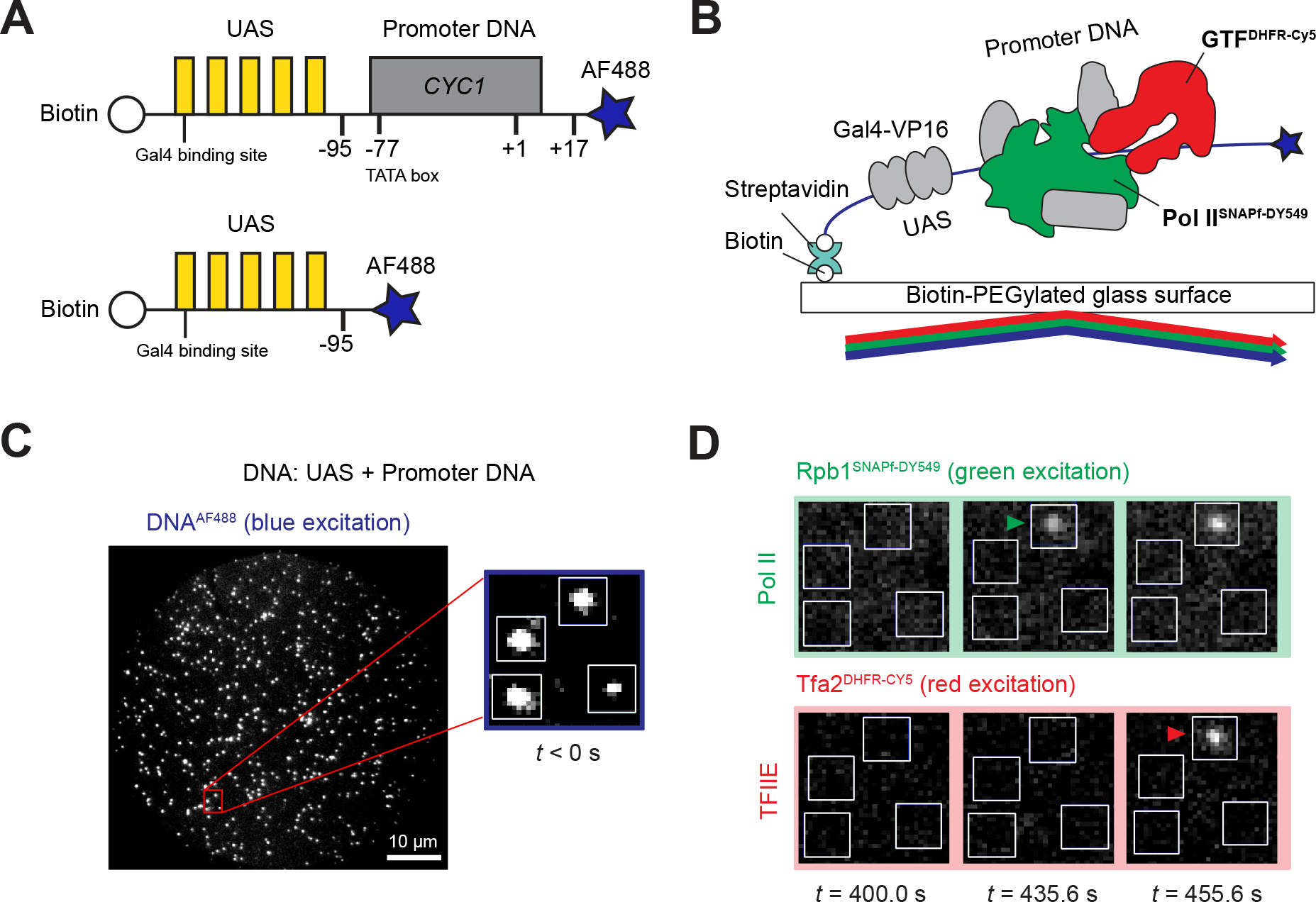
Single molecule imaging of Pol II and GTF binding to the DNA template. **(A)** Schematic of DNA templates used. UAS+promoter (*top*) has five Gal4-binding sites (yellow) upstream of the *CYC1* core promoter sequence (gray) carrying a TATA box −77) and transcription start site (+1). The UAS DNA (*bottom*) has only the Gal4-binding sites. Both are biotinylated upstream and labeled with Alexa Flour 488 downstream (blue star) relative to the promoter direction. **(B)** Schematic of single molecule imaging. DNA^AF488^ molecules were immobilized onto the passivated microscope slide surface via a biotin-streptavidin-biotin linkage and their positions mapped with blue laser excitation. Gal4-VP16 (340 nM) and yeast nuclear extract containing fluorescently labeled Pol II^SNAPf-DY549^ and GTF^DHFR-Cy5^ were added and reactions were imaged with alternating excitation with green and red lasers. **(C)** Representative image in the blue-excited channel showing hundreds of DNA^AF488^ molecules that can be simultaneously monitored in a single field of view (65 x 65 µm). Red box marks a magnified area with four DNA molecules (white spots) shown in panel D. **(D)**. Images of the red- and green-excited fluorescence channels at three time points. The indicated time is measured from the initiation of red/green imaging (*t* = 0 s), which is typically within 2-3 minutes of extract addition. Pol II and TFIIE arrivals are marked by green and red arrowheads, respectively. See also **Figure S1**.

After imaging the positions of DNA molecules, Gal4-VP16 activator (Sadowski et al., 1988) and nuclear extract were introduced into the flow chamber. Unless otherwise noted, no NTPs were added to the extract. There is no RNA synthesis under these conditions (Joo et al., 2017), allowing us to isolate and measure properties of PICs. For these experiments, ATP was further depleted with hexokinase and glucose to prevent the TFIIH translocase and kinase activities from destabilizing PICs. The arrival and departure of individual Rpb1^SNAPf-DY549^ and GTF^DHFR-Cy5^ molecules were monitored by alternating laser excitation at 633 nm (red) and 532 nm (green). Successive images in each channel were captured every 1.4 seconds (0.5 s/image for each channel, plus switching times) over a time course of 800-1200 seconds (i.e., **Fig. 1D**). Colocalization with UAS+promoter or UAS DNAs was determined using fluorescence intensity and proximity thresholds as previously described (Friedman and Gelles, 2015).

### Activator-dependent Pol II recruitment does not require the core promoter

Representative time records of Rpb1^SNAPf-DY549^ fluorescence colocalized with individual UAS+promoter DNA molecules are shown in **Fig. 2A**. As previously observed in the presence of NTPs (Rosen et al., 2020), binding events of various durations were recorded. Individual DNAs often exhibited serial Pol II binding and dissociation events (**Fig. 2A**, top). Interestingly, simultaneous binding of multiple Pol II molecules also occurred (∼45% of total Pol II binding events), seen as multiple fluorescence intensity steps (**Fig. 2A**, middle & bottom, brackets). To provide an overview of the reaction, binding records for 100 randomly chosen DNA molecules were converted to binary format indicating when at least one Pol II was present (color) or absent (white) on the DNA. The resulting horizontal time ribbons were then stacked to form a rastergram and sorted by the order of first Pol II binding from bottom to top (**Fig. 2B**, top). Most DNA templates bound Pol II at least once during the twenty minutes of imaging.

**Figure 2.**
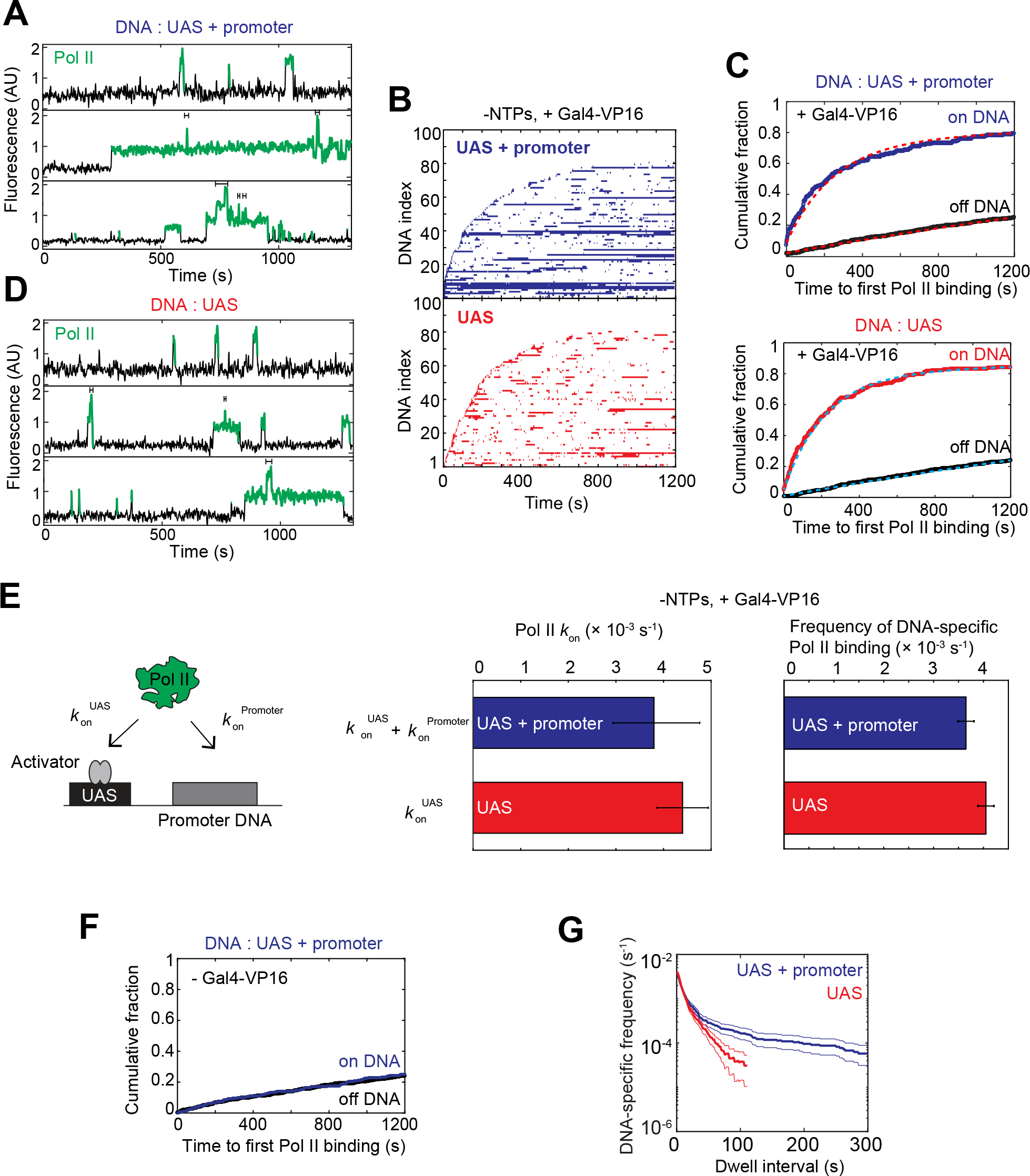
Pol II association kinetics are independent of the core promoter. **(A)** Time records of Pol II fluorescence at three different UAS+promoter locations. Green-colored intervals show times when Pol II colocalizes with the DNA template. Brackets above traces mark when multiple Pol II molecules are simultaneously bound to the same DNA. **(B)** Rastergrams show Pol II binding to 100 randomly selected DNA locations, ordered by time of first Pol II detection. Each horizontal line represents a single DNA location, and a colored band indicates one or more colocalized Pol II molecules. The *top* panel data (blue) shows UAS+promoter while the *bottom* panel data (red) shows UAS only. Note that both templates were simultaneously imaged on the same slide surface to ensure identical conditions. This reaction contained Gal4-VP16 activator, but lacked NTPs. **(C)** Cumulative fraction of DNA molecules bound at least once by Pol II as a function of time (*top*: UAS+promoter (blue), *bottom*: UAS (red)). Binding to off DNA sites (black) is shown as a control for background. Data were fit to a single-exponential specific binding model (dashed lines). Fit parameters and numbers of observations for all fits are given in panel E (center) and **Table S4**. **(D)** Time records of Pol II fluorescence at three different UAS locations, plotted as in panel A. **(E)** Schematic at left shows two pathways for initial Pol II association with the DNA template: (1) Pol II recruitment to UAS-bound activator (rate constant *k*_on_^UAS^) and (2) direct Pol II recruitment to core promoter (*k*_on_^Promoter^). Center panel shows apparent first-order rate constants (±S.E.) of Pol II initial association with the UAS+promoter (blue) or UAS (red) calculated from curve fitting in panel C. Right panel shows DNA-specific binding frequencies (±S.E.) of total (initial + subsequent) Pol II binding to Pol II-unoccupied DNA. **(F)** Cumulative fraction of Pol II-bound DNA versus time for UAS+promoter (blue) or off DNA sites (black), in the absence of Gal4-VP16. **(G)** Cumulative distribution of Pol II dwell intervals on UAS+promoter (blue) or UAS (red) with 90% confidence intervals (thin lines). Each continuous time interval with one or more labeled Pol II molecules present was scored as a single dwell. Frequency values on the vertical axis include subtraction of off DNA background. Values of zero or below after background subtraction are not plotted, which is why the UAS curve only extends to ∼100 s. See also **Figure S2**.

The cumulative fraction of UAS+promoter DNA templates bound at least once by Pol II was plotted as a function of time to first Pol II binding (**Fig. 2C**, top). Whereas 82 ± 3% (S.E.) of UAS+promoter DNA molecule sites colocalized with Pol II at least once during the 1200 s imaging interval (blue), only 25 ± 2% (S.E.) of off DNA locations did (black), demonstrating that Pol II binding was predominantly DNA-specific. The data fit well to a single-exponential specific binding model (Friedman and Gelles, 2015) (**Fig. 2C**, top, red dashed lines and **Table S4**), with initial specific binding of Pol II to DNA behaving as a single rate-limiting process with an apparent first-order association rate constant, (3.8 ± 1.0) × 10^-3^ s^-1^ (**Fig. 2E**, left bar graph, blue), similar to that measured in the presence of NTPs (Rosen et al., 2020). DNA-specific Pol II binding was completely activator-dependent, as control experiments lacking Gal4-VP16 showed no DNA binding above the non-specific background (**Fig. 2F**).

Presumably only one PIC can occupy the core promoter at a time, raising the question of how multiple polymerases can bind simultaneously (e.g. **Fig. 2A**, middle & bottom, brackets). In our earlier study, we showed that overall Pol II binding to DNA behaved as a single rate-limiting step, yet there was a time lag before binding of the subset of Pol II molecules that proceeded on to elongation. Kinetic modeling suggested the existence of an additional slow step needed to allow Pol II binding to progress to PIC and elongation complexes (Rosen et al., 2020). We postulated that Pol II might initially bind Gal4-VP16 at the UAS, with arrival of TBP or other GTFs at the core promoter being rate-limiting for subsequent Pol II incorporation into the PIC. Initial interaction with activator at the UAS could also explain simultaneous occupancy by multiple Pol II molecules. In these experiments, the concentration of Gal4-VP16 (340 nM) was in large excess over DNA molecules (<10 pM) and well above the ∼1 nM dissociation constant (Taylor et al., 1991). Therefore, each of the five Gal4 binding sites is predicted to be frequently occupied by Gal4-VP16 molecules, and multiple activators present at the UAS could tether multiple Pol II molecules.

To test this idea, we analyzed Pol II binding to a DNA template containing only the UAS in the presence of Gal4-VP16 (**Fig. 1A**, bottom). Supporting the hypothesis, UAS-associated Gal4-VP16 alone was sufficient for both initial Pol II recruitment (**Fig. 2B-C**, bottom) and simultaneous binding of multiple molecules (**Fig. 2D**, brackets). Importantly, the yeast Gcn4 activation domain (Gal4-Gcn4) was also sufficient for Pol II recruitment to the UAS (**Fig. S2A**), suggesting that Pol II recruitment to the UAS may be a general mechanism by which activators enhance PIC assembly. Pol II binding to the UAS is activator-dependent, as no binding above background was seen in the absence of activator (**Fig. S2B**). We conclude that one or more Pol II molecules, presumably complexed with Mediator, can bind UAS-bound activators independently of the core promoter.

Is this Pol II binding at the UAS on the pathway to PIC formation, or a separate, dead-end pathway? To address this question, the apparent rate constants of initial Pol II binding to UAS (*k*_on_^UAS^) and UAS+promoter (*k*_on_^UAS^ + *k*_on_^Promoter^) were compared (**Fig. 2E**, schematic). A direct promoter binding path should increase Pol II binding rate on UAS+promoter relative to UAS by *k*_on_^Promoter^. As seen for UAS+promoter, initial Pol II binding to UAS fit a single exponential model (**Fig. 2C**, bottom, cyan dashed lines and **Table S4**). The essentially identical rates of first binding at the two DNAs (**Fig. 2E**, left bar graph) suggest that any Pol II going directly to the core promoter cannot be more than a very small fraction of total binding. When we substituted the *CYC1* core promoter with the *HIS4* core promoter, Pol II was still initially recruited to the UAS and not the core promoter (**Fig. S2C-E**). The association rate constant of initial Pol II binding was identical between the *CYC1* and *HIS4* core promoters (**Fig. S2E,** right, and **Table S4**), suggesting that initial Pol II recruitment in activator-dependent transcription is independent of the core promoter. This conclusion is further supported by calculating the DNA-specific frequencies for total (rather than only initial) Pol II binding events, where UAS and UAS+promoter were again essentially the same (**Fig. 2E**, right bar graph). These results suggest that, in the context of nuclear extract, Pol II molecules predominantly bind first to UAS-bound activators on the pathway to PIC formation.

One significant difference was seen between UAS and UAS+promoter binding. A cumulative distribution of Pol II dwell times on DNA was plotted after subtracting the off DNA background values (**Fig. 2G**). The curve slopes show different Pol II dwell duration patterns on the two fragments. The nearly straight line seen on the UAS template signifies characteristically short-lived Pol II dwell times. In contrast, Pol II on UAS+promoter showed at least two different characteristic dwell times: one similar to that on UAS alone, and a longer-lived component (**Fig. 2G**). This difference is also apparent in rastergrams of the two DNAs, where many more long duration events can be seen on UAS+promoter (**Fig. 2B** and **Fig. S2A**).

Several observations suggest the long-lived Pol II dwell component results from PIC formation. In addition to dependence on core promoter, typical Pol II dwell times were strongly decreased by the inclusion of NTPs (**Fig. S2F-G**), which trigger PIC dissociation due to initiation and elongation. Furthermore, the GTFs TFIIE and TFIIF show similarly increased stable binding on UAS+promoter relative to UAS (see below). The long-lived Pol II component made up only 10-20% of total events. This suggests formation of complete PICs is relatively inefficient, but we note this number is similar to estimates of template usage for *in vitro* transcription in yeast nuclear extracts (Verdier et al., 1990; Yudkovsky et al., 2000).

Although Pol II can directly bind a TBP-TFIIB-TATA complex in the absence of activators when using purified factors (Buratowski et al., 1991; He et al., 2013; Parvin and Sharp, 1993), this does not seem to be the major pathway in the more physiological context of nuclear extract. Our results suggest that initial Pol II association with the template is primarily via transcription activators bound to the UAS/enhancer. We propose that this associated Pol II species can transfer to a PIC at the core promoter when TBP/TFIID, TFIIB, and any other required factors are present. We note that a reservoir of multiple Pol II complexes tethered at the UAS/enhancer could facilitate transcription bursts of closely spaced initiation events.

### TFIIF can bind Pol II before or after association with the template

TFIIF was first purified by its high affinity for Pol II (Sopta et al., 1985), and extensive contacts exist between TFIIF and Pol II in the PIC (He et al., 2013; Plaschka et al., 2016). However, Pol II can bind a TBP-TFIIB-TATA complex *in vitro* in the absence of TFIIF (Buratowski et al., 1991; He et al., 2013; Parvin and Sharp, 1993), and roughly half of Pol II molecules in yeast nuclear extract are not associated with TFIIF (Rani et al., 2004). Therefore, it remains unclear if Pol II incorporation into the PIC requires TFIIF pre-association, as is often assumed (Sopta et al., 1985). To determine the timing of Pol II and TFIIF recruitment, we used CoSMoS to examine their relative binding order on DNA templates.

A yeast strain was created in which Rpb1 is fused to SNAP_f_, and Tfg1, the large subunit of TFIIF, is fused to DHFR (YSB3551, see **Table S1** and **Fig. S1A, C**, and **D**). The dynamics of Rpb1^SNAPf-DY549^ and Tfg1^DHFR-Cy5^ on DNA were monitored in the same reaction by alternating red and green laser excitation, with both UAS and UAS+promoter on the same slide. Pol II and TFIIF colocalizations (i.e., time intervals during which both proteins were present) were frequently observed (**Fig. 3A**). 1475 colocalizations of Pol II and TFIIF were observed at 415 DNA locations, while only 49 colocalizations were detected at 684 off DNA locations, showing that colocalization of Pol II and TFIIF is DNA-specific. TFIIF binding to DNA templates was also highly activator-dependent (compare **Fig. 3F** and **Fig. S3A**). Like Pol II, multiple TFIIFs could simultaneously bind a single DNA (∼10% of total TFIIF binding events). Of these multiple TFIIF events, at least 80% occurred when multiple Pol II molecules were also present (brackets in **Fig. 3A** and **S3C**).

**Figure 3.**
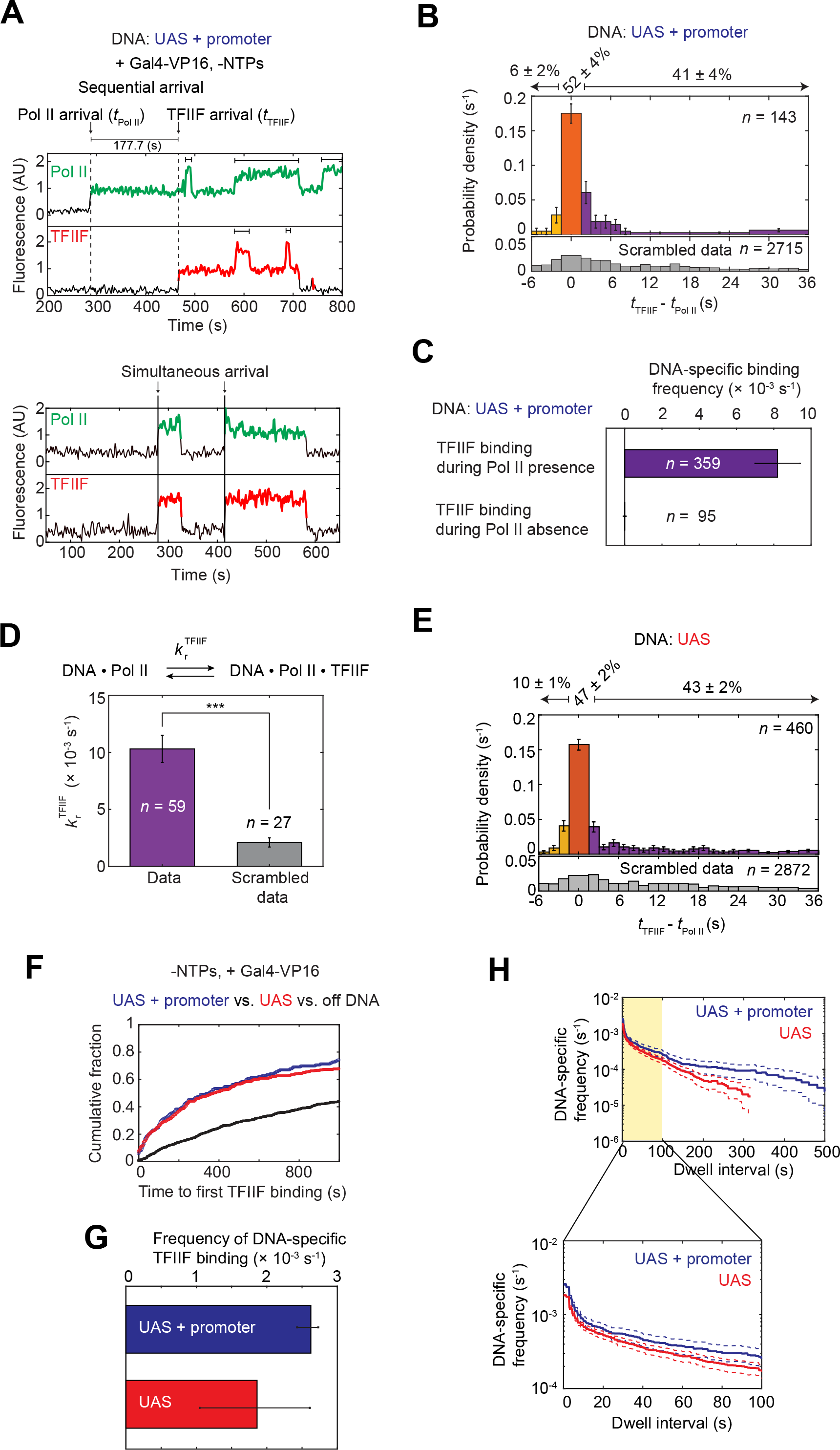
Dynamics of Pol II and TFIIF during activator-dependent PIC assembly. **(A)** Representative time records of Pol II and TFIIF at two UAS+promoter DNA molecules. Colored intervals are times when Pol II (green) or TFIIF (red) colocalized with DNA. Arrows mark arrival times of Pol II (*t*_Pol II_) or TFIIF (*t*_TFIIF_). Dashed lines on the *top* record show the sequential arrival of Pol II and then TFIIF. Solid lines on the *bottom* record show two simultaneous arrivals of Pol II and TFIIF. Brackets above traces indicate when multiple Pol II or TFIIF molecules were simultaneously bound to the same DNA. **(B)** Histogram shows probability density (±S.E.) for time differences between TFIIF and Pol II arrival (*t*_TFIIF_ − *t*_Pol II_) at unoccupied UAS+promoter DNA. Note that while 85% of *t*_TFIIF_ − *t*_Pol II_ values occur in the range shown, the percentage numbers atop the plot refer to all events. Orange bar represents apparently simultaneous Pol II and TFIIF arrival times, i.e. within ±1 video frame (±1.4 s). Purple bars represent Pol II arrivals before TFIIF, and yellow bars TFIIF arrivals before Pol II. The *bottom* histogram (gray) shows a control analysis using data scrambled by random pairing of time records. *n* = total number of the *t*_TFIIF_ − *t*_Pol II_ values. **(C)** Bars show DNA-specific binding frequencies (± S.E.) of TFIIF during time intervals when Pol II was present (purple) or absent (yellow) on UAS+promoter templates. *n* specifies the total number of TFIIF binding events within each category. **(D)** Recruitment rate constants (± S.E.) for TFIIF by Pol II (purple) or in scrambled data (gray). Asterisks indicate *P* < 0.001, calculated using paired t-test. **(E)** Histogram shows the probability density (± S.E.) for *t*_TFIIF_ − *t*_Pol II_ values at unoccupied UAS template, plotted as in panel B (81% of the *t*_TFIIF_ − *t*_Pol II_ values are shown). **(F)** Cumulative fractions over time for UAS+promoter (blue), UAS (red), and off DNA sites (black) bound at least once by TFIIF. **(G)** DNA-specific binding frequencies (±S.E.) of total TFIIF binding events to UAS+promoter (blue) or UAS (red). **(H)** Cumulative distribution of TFIIF dwell intervals on UAS+promoter (blue) or UAS (red) with 90% confidence intervals (dashed lines). Frequency values on the vertical axis are after subtraction of off DNA background. Values of zero or below after background subtraction are not plotted. *Bottom*: magnified view of the first 100 s (yellow). **(A-H)** All data were taken from an experiment using a Rpb1^SNAPf-DY549^/Tfg1^DHFR-Cy5^ yeast nuclear extract from YSB3551 (**Table S1**). See also **Figure S3**.

The binding order of Pol II and TFIIF at individual UAS+promoter DNA molecules was assessed by measuring the intervals between their arrival times (*t*_TFIIF_ –*t*_Pol II_, as diagrammed in **Fig. 3A**, top panel). For this analysis, only de novo colocalization events (i.e., starting with DNA unoccupied by either factor) were included. If the time difference was within the imaging time resolution (± 1 video frame, i.e. ± ∼1.4 s), the corresponding colocalization was scored as an apparent simultaneous arrival (**Fig. 3B**, orange bar). A large fraction of colocalizations (52 ± 4%) showed apparent simultaneous Pol II and TFIIF arrival (i.e., **Fig. 3A**, bottom). In computational simulations where individual TFIIF time records were randomly paired with Pol II time records from different DNA sites (Scrambled data), coincidental Pol II and TFIIF binding was ∼7% of colocalization events, far below the observed fraction. Therefore, Pol II and TFIIF frequently bind as a preassembled complex or bind in such rapid succession that the separate binding events cannot be resolved in the experiment.

The next largest fraction of colocalizations (41 ± 4%) exhibited sequential arrival of Pol II and then TFIIF (**Fig. 3B**, purple bars). Most intervals were only a few seconds, but some were on the order of minutes (**Fig. 3A**, top). Although Cy5-TMP dye binds the DHFR tag non-covalently, the fast and tight interaction of TMP ligands with DHFR (Calloway et al., 2007) makes it unlikely that the TFIIF lag is due to reversible labeling. The non-simultaneous arrival events were also not due to non-specific DNA binding, as TFIIF binding to DNA occurred essentially only when Pol II was present (**Fig. 3C**). To confirm that the binding of TFIIF is associated with the prior presence of Pol II in the sequential arrival events, we calculated the TFIIF recruitment rate constant *k*_r_^TFIIF^ (see STAR Methods). This calculated value of *k*_r_^TFIIF^ ((10.3 ± 1.2) × 10^-3^ s^-1^) was significantly greater than that obtained from a similar analysis of the scrambled data control ((2.1 ± 0.4) × 10^-3^ s^-1^) (**Fig. 3D**). Taken together, the **Fig. 3A-D** demonstrate that TFIIF is primarily recruited to DNA in association with Pol II, but can also arrive after Pol II binding to the template.

Given that Pol II primarily binds first at the UAS, colocalizations of Pol II and TFIIF on UAS and UAS+promoter DNAs were analyzed to determine if TFIIF binding requires the core promoter. Both simultaneous and sequential arrival of Pol II and TFIIF were observed on UAS (**Fig. 3E** and **S3B**), and the relative fractions of these two pathways were similar to UAS+promoter (**Fig. 3B** and **3E**). There was little difference in the kinetics of initial TFIIF association (**Fig. 3F**, **S3D** and **Table S4**). When both initial and subsequent TFIIF binding events were analyzed, association rates were also similar or perhaps slightly increased at UAS+promoter versus UAS alone (**Fig. 3G**, see also the *y*-intercepts of survival frequency plots of total TFIIF binding in **Fig. 3H**). These results suggest that the majority of Pol II·TFIIF complexes either arrive pre-assembled or form on the UAS. However, some TFIIF molecules may go directly to the core promoter, perhaps joining the PIC after a previous TFIIF molecule has dissociated. Consistent with stable PIC formation on the core promoter, longer dwell intervals in which at least one TFIIF molecule bound were significantly more frequent on UAS+promoter than UAS (**Fig. 3H**).

### TFIIE can join Pol II-TFIIF at the UAS or core promoter

Based on gel shift experiments (Inostroza et al., 1991; Peterson et al., 1991), the prevailing stepwise assembly model proposes that TFIIE joins the PIC after both Pol II and TFIIF are bound. Yet a reason for TFIIF-dependence is not obvious, as TFIIE has far more extensive contact with Pol II than TFIIF (He et al., 2013; Schilbach et al., 2017). We considered whether TFIIF may not be essential for initial TFIIE recruitment, but is needed to stabilize its association enough to survive native gel electrophoresis. To determine the order of factor interactions in our system, TFIIE was imaged in combination with Pol II or TFIIF. To visualize Pol II and TFIIE, nuclear extract was prepared from yeast expressing Rpb1^SNAPf^ and Tfa2, the small subunit of TFIIE, fused to the DHFR tag (YSB3474, see **Table S1** and **Fig. S1A, C and D**). For simultaneous imaging of TFIIF and TFIIE, extract was prepared with Tfg1-SNAP_f_ and Tfa2-DHFR (YSB3553, see **Table S1** and **Fig. S1B-D**). As above, reactions were performed with both the UAS and UAS+promoter templates on the same slide surface, in the absence of NTPs.

Frequent colocalization of TFIIE was seen with both Pol II (**Fig. 4A**) and TFIIF (**Fig. 4B**) at both UAS+promoter (top panels) and UAS (bottom panels). Like Pol II and TFIIF, TFIIE binding was far more frequent at DNA sites compared to off DNA sites (**Fig. 4D**), and was dependent on Gal4-VP16 activator (**Fig. S4A**). Simultaneous binding of multiple TFIIE molecules was sometimes observed during periods when multiple Pol II molecules were bound (**Fig. S4C**). TFIIE bound DNA 24-fold more frequently when Pol II was present than absent, and 26-fold more frequently when TFIIF was present (**Fig. 4C**), supporting the model that both Pol II and TFIIF are needed for TFIIE recruitment. This dependence suggests that TFIIF binding triggers a conformation change in Pol II or TFIIE that allows the two to stably interact, or that TFIIE binding to Pol II without TFIIF is so unstable that dwell times are significantly below the single frame imaging time (0.5 s). The association rate constants of initial TFIIE binding were the same whether or not the DNA contained a core promoter (**Fig. 4D**, **Fig. S4B** and **Table S4**). Therefore, both TFIIE and TFIIF can associate with Pol II at the UAS, independently of a TATA box. As seen for Pol II (**Fig. 2G**), TFIIE dwell times on the UAS were short, while longer dwells, presumably in the PIC, were seen on UAS+promoter (**Fig. 4E**).

**Figure 4.**
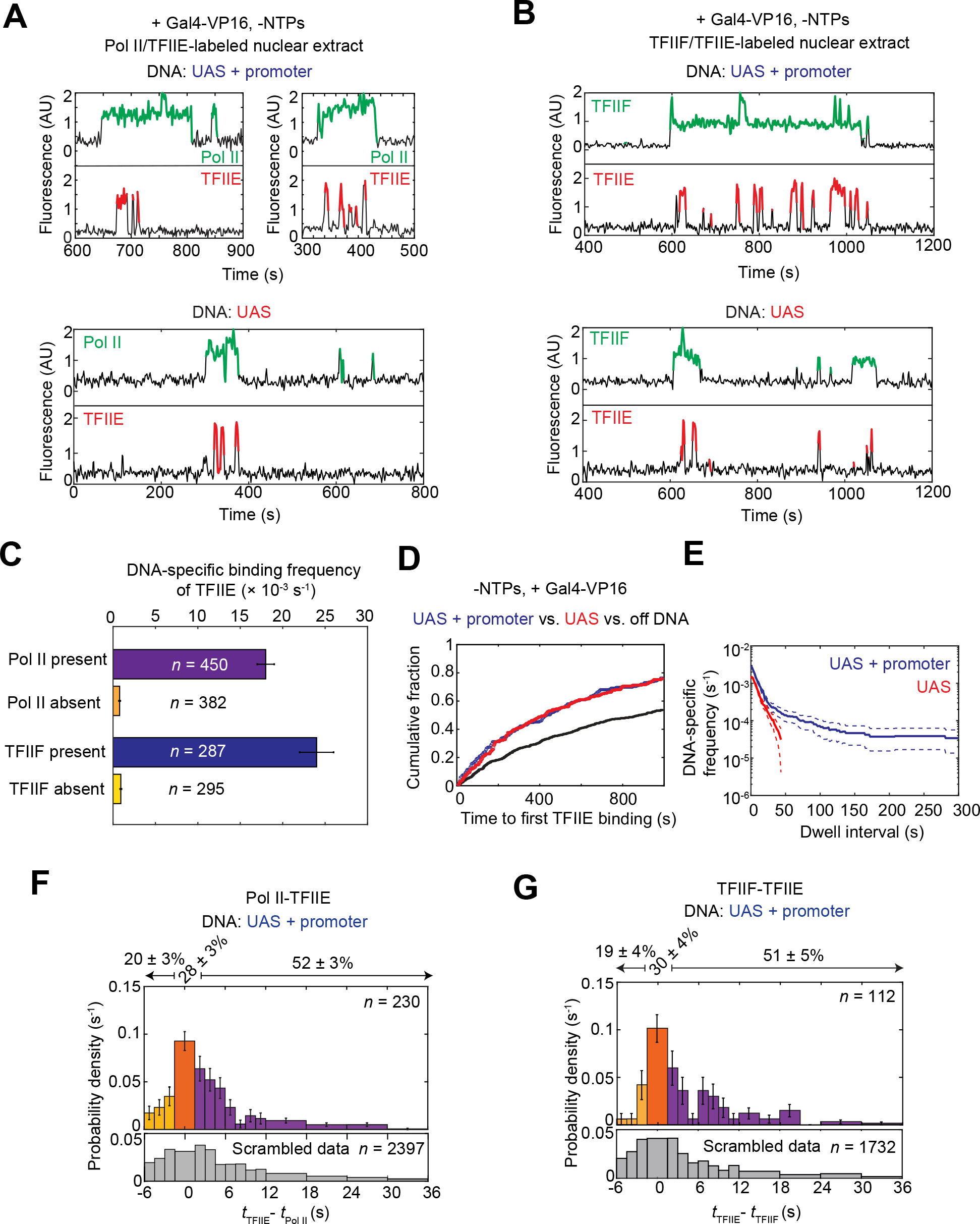
Dynamics of TFIIE relative to Pol II and TFIIF during activator-dependent PIC assembly. **(A)** Representative time records at two UAS+promoter molecules (*top*) and one UAS molecule (*bottom*) for Pol II (green) and TFIIE (red) colocalization with DNA. **(B)** Time records at one UAS+promoter molecule(*top*) and one UAS molecule (*bottom*) for TFIIF (green) and TFIIE (red) colocalization with DNA. **(C)** DNA-specific binding frequencies (±S.E.) of TFIIE during time intervals when Pol II was present (purple) or absent (orange), and when TFIIF was present (blue) or absent (yellow) at UAS+promoter. *n* = number of TFIIE binding events used in each calculation. **(D)** Cumulative distributions over time for the fractions of the UAS+promoter (blue), UAS (red) and off DNA sites (black) bound at least once by TFIIE. **(E)** Cumulative distribution of TFIIE dwell intervals on UAS+promoter (blue) or UAS (red), with 90% confidence intervals (dashed lines). Frequency values are after subtraction of off DNA background; values of zero or below are not plotted. **(F)** Histogram of probability density (±S.E.) for time differences between TFIIE and Pol II arrival times (*t*_TFIIE_-*t*_Pol II_) at unoccupied UAS+promoter DNA, plotted as in **Fig. 3B** (83% of the *t*_TFIIE_-*t*_Pol II_ values are within shown range). **(G)** Histogram of probability density (±S.E.) for time differences between TFIIE and TFIIF arrival times (*t*_TFIIE_ –*t*_TFIIF_) at occupied UAS (80% of the *t*_TFIIE_-*t*_TFIIF_ values shown). **(A-G)** Simultaneous Pol II and TFIIE fluorescence imaging was done using Rpb1^SNAPf-DY549^/Tfa2^DHFR-Cy5^ nuclear extract from YSB3474, while TFIIF and TFIIE co-imaging used Tfg1^SNAPf-DY549^/Tfa2^DHFR-Cy5^ nuclear extract from YSB3553 (**Table S1**). Data in panels D and E is from YSB3474. See also **Figures S4** and **S5**.

TFIIE binding showed one striking difference from TFIIF (**Fig. 3A**) and TFIIH (see below), in that multiple cycles of TFIIE association and dissociation were often seen during each TFIIF or Pol II binding event (**Fig. 4A** and **4B**). This repetitive binding also manifests in the total TFIIE binding frequency (**Fig. S5A**), where UAS+promoter had a higher frequency ((2.87 ± 0.15) × 10^-3^ s^-1^) than UAS ((1.49 ± 0.11) × 10^-3^ s^-1^). Repetitive TFIIE binding is more likely during long duration Pol II or TFIIF binding, and is therefore preferentially seen on UAS+promoter DNA. These results suggest a model where TFIIE first joins the Pol II·TFIIF complex primarily at the UAS, but also frequently exchanges on and off Pol II·TFIIF complexes that have transferred to the core promoter.

To determine if TFIIE pre-binds Pol II or TFIIF before arrival at the template, time differences between Pol II and TFIIE arrival (*t*_TFIIE_–*t*_Pol II_) (**Fig. 4F**), and between TFIIF and TFIIE (*t*_TFIIE_–*t*_TFIIF_) (**Fig. 4G**) were calculated. The small fraction of TFIIE arrivals before Pol II or TFIIF (yellow bars in **Fig. 4F** and **4G**) may be coincidental independent binding, as frequencies were similar to those calculated using scrambled control data (**Fig. S5B**, yellow bars). In roughly half the co-binding events on DNA, TFIIE arrived after Pol II or TFIIF (purple bars in **Fig. 4F** and **4G**). An additional ∼30% of co-binding events were scored as apparently simultaneous within the imaging time resolution (orange bars in **Fig. 4F** and **4G**). However, when the sequential binding data (**Fig. 4F**, purple bars) were fit to a first-order sequential binding model curve for Pol II and TFIIE (**Fig. S5C**) and extrapolated back to time 0, ∼60% of the apparently simultaneous binding could be accounted for by sequential binding of TFIIE in the time interval 0 ≤ *t*_TFIIE_ –*t*_Pol II_ <1.4 seconds. We therefore conclude that TFIIE is unlikely to appreciably preassemble with Pol II or TFIIF in yeast nuclear extract.

If TFIIE is unlikely to pre-bind Pol II·TFIIF complexes off the DNA, what triggers their association on the template? Binding occurs on both UAS and UAS+promoter, with no change in the binding order or kinetics (**Fig. S5D**). Therefore, TFIIE binding is not dependent on Pol II·TFIIF interactions with factors at the TATA box. TFIIE binding is present for ∼19% of the time that Pol II or TFIIF is bound to the UAS template. If these factor interactions on DNA, presumably occurring while tethered to Gal4-VP16, were similar in solution, we would predict an equal fraction of simultaneous arrivals rather than the calculated fraction of Pol II and TFIIE simultaneous arrivals (9 ± 5%) (**Fig. S5C**). This difference suggests that activator interaction with Pol II·TFIIF might somehow promote TFIIE binding. We speculate this could involve activator-induced conformation changes transmitted via Mediator, which links activation domains and Pol II·TFIIF.

### TFIIH requires the core promoter DNA for PIC incorporation

To examine if initial TFIIH association with DNA is also independent of core promoter as seen for Pol II, TFIIF and TFIIE, we monitored Pol II and TFIIH using fluorescently labeled nuclear extract containing Rpb1^SNAPf-DY549^ and the tagged TFIIH subunit Tfb1^DHFR-Cy5^ (**Fig. S1** and **Table S1**). Like the other GTFs tested, DNA-specific TFIIH binding was activator-dependent (compare **Fig. 5A** and **S6A**). However, in marked contrast to Pol II, TFIIF, and TFIIE, binding of TFIIH above background was seen only on the UAS+promoter template, not UAS only (**Fig. 5A**). TFIIH dwell times in the absence of NTPs tend to be long, similar to the other GTFs at the core promoter (**Fig. 5B**). Strikingly, only one molecule of TFIIH associates at a time (**Fig. S6D**). Whereas multiple Pol II, TFIIF, and TFIIE molecules can associate with templates via the five activator-bound Gal4 sites and the core promoter, TFIIH apparently only binds the single PIC expected on the core promoter.

**Figure 5.**
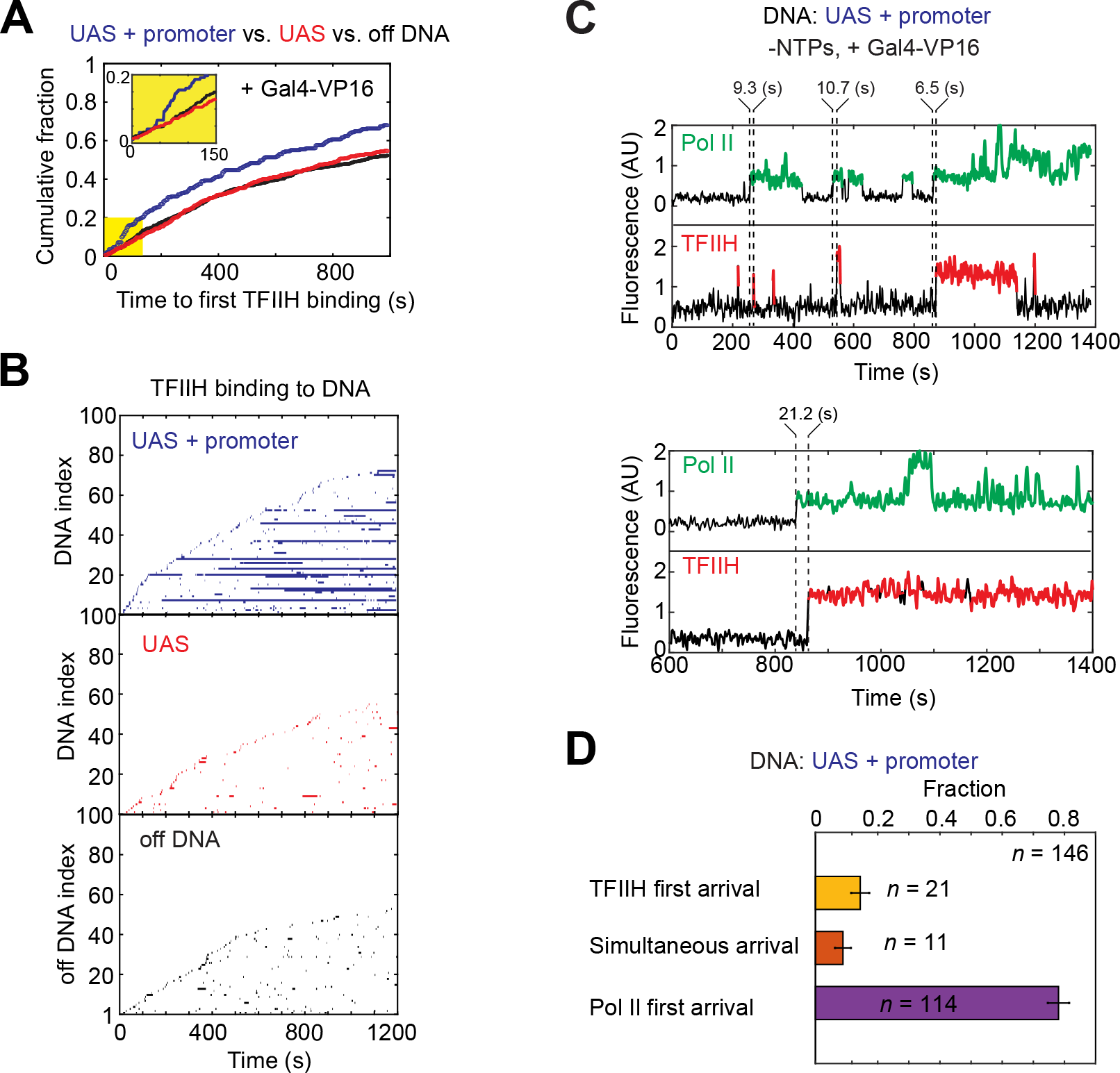
Dynamics of TFIIH relative to Pol II during activator-dependent PIC assembly. **(A)** Cumulative fraction versus time for UAS+promoter (blue), UAS (red), and off DNA (black) locations bound at least once by TFIIH. Inset: magnified view of the first 150 s (yellow) showing lag in TFIIH binding. **(B**) Rastergrams of TFIIH dwells at 100 randomly selected on or off DNA locations, plotted as in Fig. 2B. UAS+promoter (*top*, blue), UAS only (*middle*, red), Off DNA (*bottom*, black). **(C)** Representative time records of Pol II (green) and TFIIH (red) colocalizations at two UAS+promoter molecules. Dashed lines indicate the arrival times of Pol II or TFIIH, with time differences shown above. **(D)** Fractions (± S.E.) of the formation pathways for Pol II·TFIIH complexes on UAS+promoter DNAs. See **Fig. S6B** for time difference histogram. **(A-D)** Data were taken from an experiment using NTP-depleted Rpb1^SNAPf-DY549^/Tfb1^DHFR-Cy5^ nuclear extract from YSB3473 (**Table S1**). See also **Figure S6**.

Importantly, the cumulative fraction plot of times to first TFIIH binding at each DNA template revealed a non-exponential distribution, with a pronounced lag of ∼50 s after the start of imaging (**Fig. 5A**, inset). The simplest explanation for the TFIIH lag is that slower binding of one or more others factors must occur first. This slow step is likely to be required for transfer of Pol II and associated factors from the UAS to the core promoter. Notably, Spt5 association shows a similar lag under transcription conditions (Rosen et al., 2020). TFIIH almost exclusively arrived after Pol II (78 ± 3 (S.E.)% of colocalizations), confirming sequential binding (**Fig. 5C, 5D**, and **S6B**). Also consistent with a late step in PIC assembly, the delay time between TFIIH and Pol II arrival was significantly longer than that between Pol II and TFIIE (**Fig. S6C**). We thus conclude that TFIIH binding occurs after transfer of Pol II (likely with TFIIF and TFIIE) from the UAS to the core promoter.

## DISCUSSION

Under physiological conditions, numerous proteins are needed to form Pol II PICs. These include GTFs, transcription activators, and coactivators. Yet current PIC assembly models are based primarily on activator-independent studies using only purified GTFs. These experiments led to a simple sequential binding pathway directly on the core promoter. While not excluding this model, our single molecule experiments reveal that a more complex, activator-dependent pathway predominates in the more physiological context of nuclear extract (**Fig. 6A**). Surprisingly, initial template association of Pol II, TFIIF, and TFIIE requires only the UAS and activator, independent of the core promoter. However, these initial UAS-bound complexes are relatively short-lived, on the order of a few seconds. The presence of core promoter sequences produces longer duration complexes that exhibit properties expected of PICs, such as sensitivity to NTPs. The similar initial association rates observed on UAS versus UAS+promoter are most simply explained by intermediate complexes assembled on the UAS before transfer to the core promoter (**Fig. 6A**, blue box). Such intermediates also fit a kinetic model derived from our earlier study imaging Pol II and the elongation factor Spt4/5 (Rosen et al., 2020). It is notable that this branched model also fits earlier activator-independent experiments with purified GTFs. The gel shifted species that led to the classic sequential assembly pathway (Buratowski et al., 1989; Inostroza et al., 1991; Peterson et al., 1991) are encompassed in the core-promoter complexes (**Fig. 6A**, yellow box).

**Figure 6.**
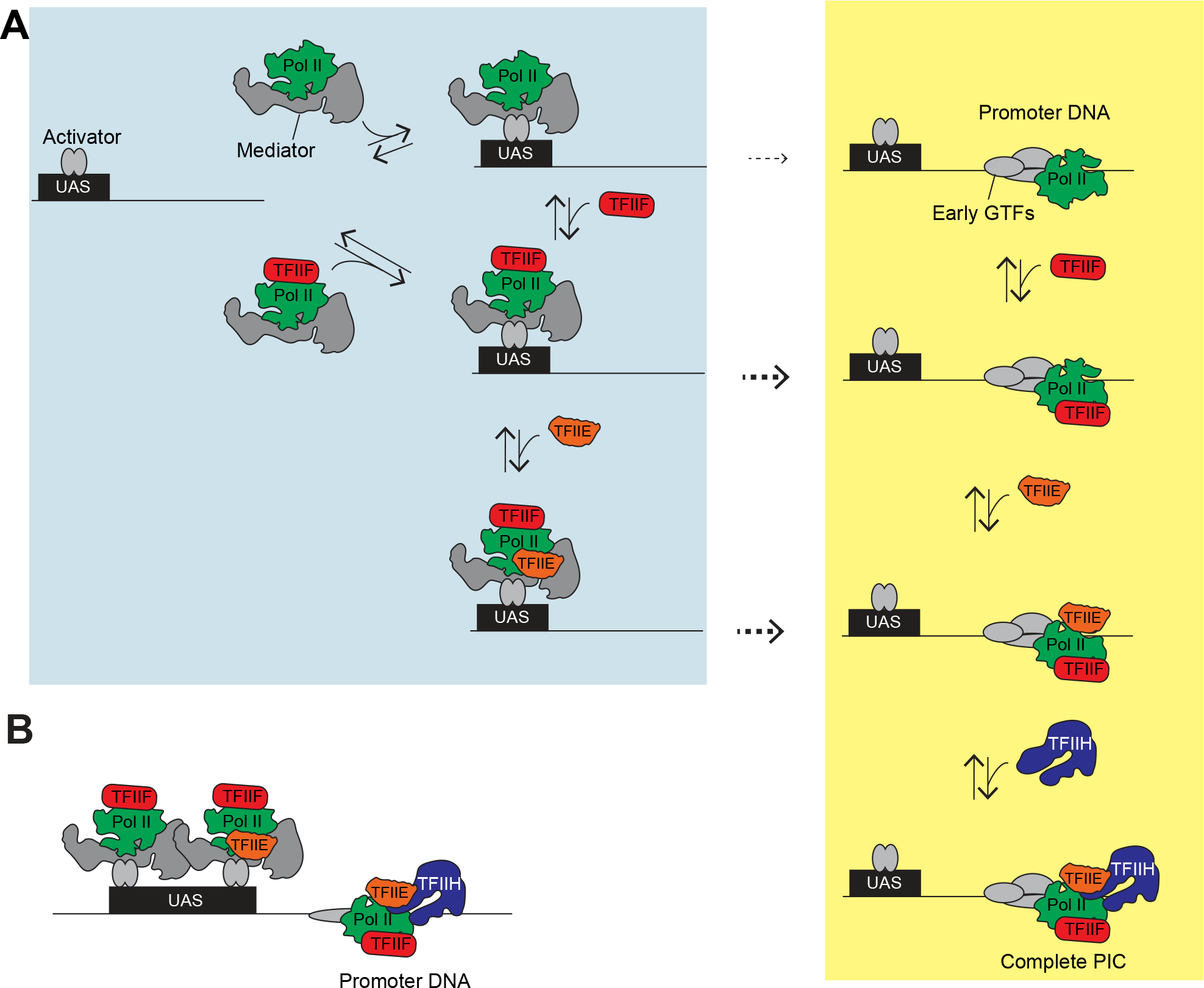
Branched model for activator-dependent PIC assembly pathways. **(A)** The single molecule imaging leads to a model where initial interactions (blue box) of Pol II (green) with the template are primarily via UAS-bound transcription activators (gray ovals), presumably via Mediator (gray), either in association with TFIIF (red) or by itself. TFIIF can also be recruited to Pol II already bound at the UAS. TFIIE (orange) is more dynamic and can join the Pol II·TFIIF complex at the UAS or core promoter. The dashed arrows indicate transfer of Pol II, Pol II·TFIIF, or Pol II·TFIIF·TFIIE from the UAS to the core promoter. TFIIH is recruited directly to the PIC at the core promoter. This model also predicts that in a non-physiological, activator-independent reaction, TFIIF, TFIIE, and TFIIH would incorporate through the intermediates at the core promoter (yellow box), thereby encompassing the previous sequential assembly model. **(B)** This activator-dependent PIC assembly model also explains the observation of simultaneous binding of multiple Pol II, TFIIF, or TFIIE molecules on the same DNA, tethered by the multiple activators at the UAS/enhancer plus the single PIC. This phenomenon could provide a mechanism for the in vivo clustering of Pol II observed by microscopy, either in cooperation with or independently of proposed condensate formation.

Could the partial PICs on the UAS correspond to previously proposed Pol II “holoenzyme” complexes? This is unlikely for several reasons. First, a substantial fraction of TFIIF and the majority of TFIIE only bind DNA after Pol II arrives (**Fig. 3E, 4F-G** and **S5D**), arguing against pre-assembly of holoenzymes off the DNA, at least in yeast nuclear extract. Second, both yeast and mammalian holoenzyme were reported to contain TFIIH (Kim et al., 1994; Koleske and Young, 1994; Maldonado et al., 1996; Ossipow et al., 1995; Ranish et al., 1999), but this factor does not stably interact with Pol II, TFIIF, and TFIIE at the UAS, and instead requires the core promoter for stable binding (**Fig. 5A** and **5B**). The very different dynamics of TFIIF, TFIIE, and TFIIH relative to Pol II and each other are inconsistent with a preassembled holoenzyme.

The association of TFIIF and TFIIE with Pol II before core promoter binding is notable from an evolutionary perspective. Subunits of RNA polymerases I and III are generally homologous or identical to those in Pol II. However, several additional Pol I and III subunits have no Pol II counterparts, but instead structurally resemble TFIIF or TFIIE (for review, see (Engel et al., 2018; Khatter et al., 2017; Vannini and Cramer, 2012)). Under particular *in vitro* conditions, TBP and TFIIB are sufficient for Pol II binding to the core promoter and accurate initiation *in vitro* (Buratowski et al., 1991; Parvin and Sharp, 1993). Although TFIIF and TFIIE are therefore not absolutely required for initial Pol II binding to the TBP-TFIIB-promoter complex, the Pol II (He et al., 2016; Murakami et al., 2015; Plaschka et al., 2016) and Pol III (Han et al., 2018) PIC structures show TFIIF or the homologous Pol III subunit contacting TFIIB or TFIIIB, respectively. This interaction, together with TFIIF and TFIIE encircling and constraining template DNA downstream of the TATA box, may further position and stabilize polymerase within the PIC. An interesting evolutionary question is whether the single RNA polymerase in eukaryotic ancestors had integral subunits that branched off in the Pol II system to become TFIIF and TFIIE, or instead began with independent initiation factors that evolved into Pol I and III subunits.

Another unexpected result from our study is that TFIIE shows repetitive cycles of binding to the PIC, perhaps explaining weaker TFIIE binding inferred from crosslinking and cryoEM studies (Grunberg et al., 2012; He et al., 2013). In the absence of NTPs to trigger promoter melting and initiation, both TFIIF and TFIIH show core promoter-dependent dwells that can last for minutes. In contrast, TFIIE rapidly exchanges on and off, even when Pol II or TFIIF is stably bound (**Fig. 4A** and **4B**). TFIIE is reportedly required for TFIIH binding (Compe et al., 2019; Maxon et al., 1994), and PIC structures show direct contacts between the two factors (He et al., 2016; Schilbach et al., 2017). Yet long dwell times of TFIIH but not TFIIE are difficult to reconcile with the classic PIC assembly model. It may be that TFIIE promotes initial TFIIH binding, but is not required to retain TFIIH once it makes contacts with Pol II and downstream promoter DNA.

Our discovery of partial PIC pre-assembly at the UAS provides important new insight into transcription activation. Activators are often said to “recruit” co-activators and basal machinery (Ptashne and Gann, 1997), but this vague term fits several non-mutually exclusive mechanisms. In the classic cooperative binding model, initial associations of activator and polymerase with DNA are independent, but subsequent contact between the two mutually stabilizes their binding. Transcription activation at many prokaryotic promoters fits this model (van Hijum et al., 2009). In this mechanism, polymerase dissociation rates of one or more intermediate pre-initiation complexes are reduced to increase transcription. In contrast, a kinetic enhancement model postulates that activator increases the initial association rate of RNA polymerase (or other basal factors) with the promoter.

Consistent with kinetic enhancement, Gal4-VP16 or Gal4-Gcn4 activator strongly accelerates Pol II association with the template, although initial binding is to the UAS/enhancer rather than the core promoter as typically pictured in most models. The observations of multiple Pol II, TFIIF, and TFIIE molecules simultaneously bound at the UAS (schematic in **Fig. 6B**, brackets in **Fig. 2D**, **S3B**, **S4C**) definitively demonstrate that activators can increase the local concentration of PIC components. This reservoir of factors can drive PIC formation, and may also contribute to transcription “bursting” if multiple Pol II·TFIIF·TFIIE complexes can rapidly and sequentially transfer to the core promoter during a window of TFIID and TFIIB binding.

The multiple Pol II molecules we observe at a single UAS are reminiscent of polymerase clusters observed in nuclei, which colocalize with Mediator (Cho et al., 2018; Sabari et al., 2018) (reviewed in (Cramer, 2019)). Although these clusters are often interpreted as condensates, our results suggest another plausible mechanism independent of phase separation. A single enhancer is comprised of multiple activator binding sites, and if each activator (which may itself have multiple activation domains) independently tethers one or more polymerases, a cluster of Pol II would appear in the microscope. Although the cluster would appear long-lived, individual Pol II molecules could exchange if they rapidly associate and dissociate from DNA-bound activator. Our single molecule studies show that average dwell times of Pol II tethered to the UAS by Gal4-VP16 are on the order of seconds, very similar to time scales seen for fluorescence recovery after photobleaching of Pol II clusters *in vivo* (Cho et al., 2018).

Our system for visualizing individual transcription factors in nuclear extract has revealed new intermediates not seen with purified factors, indicating that the activator-dependent pathways by which PICs form are more complicated than previously proposed. Future studies with additional labeled factors are likely to provide further new insights into Pol II transcription activation, initiation, and elongation.

### Limitations

Like other in vitro transcription systems, the factor concentrations in the extracts used here are lower than those in living cells. Therefore, the parameters measured here are “apparent” association and dissociation rates specific to these conditions. However, the relative ratios of factors and templates appear to be preserved in the reactions, so the relative orders of competing PIC assembly pathways should be similar to those in vivo. Also, the experiments here use naked DNA templates rather than chromatin, and therefore measure events that would occur after removal or remodeling of nucleosomes that occlude the promoter in vivo.

## ACKNOWLEDGMENTS

We thank members of the Buratowski and Gelles labs for helpful discussions, and Yujin Chun for technical support. This work was supported by NIH grants R01GM046498 to S.B., R01GM081648 to J.G., and R01CA246500 to S.B. and J.G. I.B. was partially supported by the Van Maanen Fellowship from Harvard Medical School.

## AUTHOR CONTRIBUTIONS

I.B. performed all experiments, and all authors contributed to designing and interpreting experiments and writing of the paper.

## STAR METHODS

### RESOURCE AVAILABILITY

#### Lead contact

Further information and requests for resources and reagents should be directed to and will be fulfilled by the Lead Contact, Stephen Buratowski.

#### Materials Availability

*S. cerevisiae* strains and plasmids generated in this study (**Table S1** and **S2**) are available from the Lead Contact upon request.

#### Data and Code Availability

• Original/source data for single molecule experiments are provided as “intervals” files. The data files have been deposited at Zenodo and are publicly available as of the date of publication. A DOI is listed in the key resources table.
• “Intervals” files can be read and manipulated by the Matlab program “Imscroll”, which is available at https://github.com/gelles-brandeis/CoSMoS_Analysis.
• Any additional information required to reanalyze the data reported in this paper is available from the lead contact upon request.

### EXPERIMENTAL MODEL AND SUBJECT DETAILS

*S. cerevisiae* strain information is in **Table S1**. All *S. cerevisiae* strains for nuclear extract preparation were grown in YPD (Yeast extract Peptone 3% Dextrose) media at 30°C until OD_600_ reached 3-4.

### METHODS DETAILS

#### Generation of HA_3_ -DHFR/SNAP_f_ tagging plasmids

A hygromycin resistant DHFR tag which also encodes for an HA tag was created by an isothermal assembly reaction with the following four DNA fragments: (1) pBlueScript (Stratagene, #212205; see **Table S2**) plasmid digested with EcoRI and BamHI, (2) a DNA fragment containing a HA tag and GSG linker repeats amplified from pFA6a-HA-KIURA3 with primers 3XHA-Forward and 3XHA-Reverse, (3) a DHFR tag amplified from pAAH2 with primers DHFR-Forward and DHFR-Reverse and (4) a hygromycin resistance cassette amplified from pAAH2 with primers TEFpro-Forward and TEFterm-Reverse. It should be noted that a DHFR tag is followed by a 39 bp synthetic terminator which prevents transcription read-through toward the hygromycin resistance cassette. The resulting plasmid pBS-SKII-3XHA-eDHFR-Hygromycin (YV317) was confirmed by sequencing.

A nourseothricin resistant fast SNAP (SNAP_f_) tag which also encodes for an HA tag was created by an isothermal assembly reaction with the following four DNA fragments: (1) pBlueScript plasmid digested with EcoRI and BamHI, (2) a DNA fragment containing a HA tag and GSG linker repeats amplified from pFA6a-HA-KIURA3 with primers 3XHA-Forward and 3XHA-Reverse, (3) a SNAP_f_ tag amplified from pAAH13 with primers SC-Forward and SC-Reverse and (4) a nourseothricin resistance cassette amplified from pAAH6 with primers TEFpro-Forward and TEFterm-Reverse. It should be noted that a SNAP_f_ tag is followed by a 39 bp synthetic terminator which prevents transcription read-through toward the nourseothricin resistance cassette. The resulting plasmid pBS-SKII-3XHA-fSNAP-NAT (YV309) was confirmed by sequencing.

Plasmid information is given in **Table S2**. Primers used to generate the plasmids are listed in **Table S3**.

#### Yeast strains, plasmids, and oligonucleotides

*S. cerevisiae* strains used in this study are listed in **Table S1**. To create doubly-fused yeast strains (YSB3473, YSB3474, YSB3551 and YSB3553 in **Table S1**), two DNA cassettes for yeast transformation were prepared by PCR amplification of YV317 (**Table S2**) and YV309 (**Table S2**) for DHFR and SNAP_f_ tagging, respectively, with the appropriate primer pairs from **Table S3**. The amplified DHFR- and SNAP_f_-containing fragments were sequentially transformed into the protease-deficient strain, YF702 (**Table S1**). After each round of transformation, positive clones were selected for the marker. Additionally, selected clones were checked for presence of the insert by colony PCR with the appropriate primer pairs from **Table S3**. In-frame fusion protein expression and stability was confirmed by immunoblotting for the target protein. The fusion strains had similar growth to the YF702 wild type as determined by a spotting assay (**Fig. S1C**).

#### Yeast nuclear extract preparation

Yeast nuclear extracts were prepared as previously described (Rosen et al., 2020; Sikorski et al., 2012). Briefly, yeast cells were grown in 4 liters of YPD (3% dextrose) medium at 30°C to an OD_600_ of 3–4 and were harvested. Yeast cell walls were digested with 15 mg of Zymolyase 100T (Amsbio, #120493-1) until ∼80%–90% of cells became spheroplasts. The digestion times varied from 30 min to 1.5 hrs depending on the strain. The spheroplasts were suspended in 250 mL of YPD+1 M sorbitol and incubated at 30°C for 30 min for recovery. After residual Zymolyase 100T was removed by serial centrifugations, the spheroplasts were resuspended in Buffer A (lysis buffer; 18% (w/v) Ficoll 400, 10 mM Tris-HCl pH 7.5, 20 mM potassium acetate, 5 mM magnesium acetate, 1 mM EDTA (ethylenediamine tetraacetic acid), 0.5 mM spermidine, 0.15 mM spermine, 3 mM DTT, 1 µg/mL each of aprotinin, leupeptin, pepstatin A, and antipain). The resuspended spheroplasts were lysed with a motorized homogenizer (Wheaton, #62400-802) and the supernatant was collected by four sequential centrifugations (twice at 5000 x *g* for 8 min followed by twice at 5000 x *g* for 5 min). Crude nuclei from the lysed spheroplasts were pelleted by centrifugation at 25,000 x *g* for 30 min and were suspended in Buffer B (100 mM Tris-acetate pH 7.9, 50 mM potassium acetate, 10 mM magnesium sulfate, 10% glycerol, 3 mM DTT, 2 mM EDTA, 1 µg/mL each of aprotinin, leupeptin, pepstatin A, and antipain). Nuclear proteins were extracted by the addition of ammonium sulfate solution (pH 7.5) to a final concentration of 0.4 M, followed by rotating for 30 min at 4°C. After centrifugation in a Beckman type 71 rotor at 37,500 rpm (∼120,000 x *g*) for 1.5 hrs at 4°C, the soluble fraction was collected. Nuclear proteins were precipitated by the addition of solid ammonium sulfate granules to ∼70% saturation (0.35 g to 1 mL of nuclear protein solution). The ammonium sulfate precipitate was recovered by centrifugations at 13,000 rpm (∼16,000 x *g*) for 20 min and then 5 min. The pooled protein pellet was weighed and resuspended in Buffer C’ (20 mM HEPES pH 7.6, 10 mM MgSO_4_, 1 mM EGTA (ethylene glycol-bis(β-aminoethylether) tetraacetic acid), 10% glycerol, 3 mM DTT, 1 µg/mL each of aprotinin, leupeptin, pepstatin A, and antipain) at a ratio of ∼400 mg protein pellet to a 1 mL Buffer C’.

SNAP_f_ fusion proteins in nuclear extracts were labeled by adding SNAP-Surface 549 (New England BioLabs, #S9112S) to a final concentration of 0.4 µM (unless otherwise specified) and incubating at 4°C for 1 hr on a rotator in the dark. The suspension was dialyzed three times (1 hr, 1.5 hrs and 2 hrs) each time against 500 mL of Buffer C’ supplemented with 75 mM ammonium sulfate (nuclear extract dialysis buffer). Residual unreacted SNAP-Surface 549 was removed from SNAP-labeled extracts as previously described (Haraszti and Braun, 2020; Rosen et al., 2020). Briefly, one fourth of the extract volume of SNAP_f_-coupled agarose beads were added to the extract and incubated at 4°C for 1 hr on a rotator, after which the beads were removed by centrifugation at 1,000 x *g* for 2 min at 4°C. Finally, the extract was aliquoted, frozen in liquid nitrogen, and stored at −80°C. Labeling of SNAP_f_ fusion proteins and depletion of residual dye were confirmed by in-gel fluorescence imaging on a Typhoon imager (GE Healthcare). *In vitro* transcription activity of the final nuclear extract was measured by the *in vitro* transcription assay described below.

#### Preparation of SNAP_f_-coupled agarose beads

His-SNAP_f_ proteins were prepared as previously described (Rosen et al., 2020). NHS-activated agarose beads (Thermofisher Scientific #26196) were incubated with purified His-SNAP_f_ protein at a ratio of 90 mg resin to 10 mg protein for 2-3 hr at room temperature. The beads were transferred to a 5 mL Pierce centrifuge column (Thermofisher Scientific, #89897). After washing with 1X PBS (137 mM NaCl, 2.7 mM KCl, 10 mM Na_2_HPO_4_, and 1.8 mM KH_2_PO_4_, pH 7.4) until residual uncoupled His-SNAP_f_ protein was no longer detectable in the effluent using Protein Detection Reagent (Pierce PI23200), resin was quenched with 1 M Tris-HCl, pH 7.5 for 30 min at room temperature. The beads were equilibrated with nuclear extract dialysis buffer supplemented with 3 mM DTT for long-term storage at 4°C. Prior to use, efficiency of the dye depletion by the beads was tested. SNAP-Surface 549 (New England Biolabs, #S9112S) was suspended in yeast nuclear extract containing no SNAP_f_ -fusion proteins to a final concentration of 1 µM. The extent of a decrease in the dye concentration after dye depletion was measured by in-gel fluorescence imaging on a Typhoon imager (GE Healthcare). The residual dye concentration was determined by comparison against a standard curve of known free dye concentrations. The SNAP_f_-coupled bead treatment typically reduced SNAP-Surface 549 concentration in extract from 1 µM to 50 nM.

#### *In vitro* bulk transcription assay

*In vitro* transcription assay was performed as previously described (Sikorski et al., 2012). The plasmid pUC18-G5CYC1 G- (SB649) (Joo et al., 2019) was used as the DNA template. The DNA template (100 ng) was incubated with Gal4-VP16 (340 nM), the ATP regeneration system consisting of creatine kinase (0.1-0.2 units) and phosphocreatine (10 mM), and nuclear extract (typically 8 µL to 10 µL in 50 µL reaction). 400 µM each of ATP, CTP, UTP and 100 µM of 3’-O-methyl-GTP (chain terminator) along with ^32^P-labeled UTP were added to initiate the reaction. After 45 min reaction at room temperature, transcripts were treated with RNAse T1 and Proteinase K, extracted with phenol-chloroform, ethanol precipitated, separated by gel electrophoresis (8 M urea 5% polyacrylamide gel), and analyzed by autoradiography and/or phosphorimager (GE Healthcare).

#### Preparation of DNA templates for single molecule assays

Upstream biotinylated and downstream AF488 labeled DNA templates (**Fig. 1A**) were prepared by PCR from pUC18-G5CYC1 G- (SB649; **Table S2**) with Platinum Taq DNA polymerase (Invitrogen) and primers Biotin-universal and AF488-G-less-PL_REV for the UAS and AF488-Mid G-less d2 REV(+17) for the UAS+promoter templates (primer sequence information in **Table S3**). The DNA templates used to test effects of the core promoters and 4NTPs were prepared by PCR from pUC18-G5CYC1 G- (SB649; **Table S2**) and pUC18-G5-HIS4 G- (SB1964; **Table S2**) for the *CYC1* and *HIS4* core promoter, respectively with Platinum Taq DNA polymerase (Invitrogen) and primers Biotin-universal and AF488-M13 rev2 (primer sequence information in **Table S3**). The PCR product was purified using DNA SizeSelector-I SPRI magnetic beads (Aline Biosciences). The DNA template sequence information is in **Table S5**.

#### Single-molecule microscopy

Single-molecule imaging experiments were performed on a multi-wavelength single-molecule total internal reflection fluorescence (TIRF) inverted microscope with a pair of micromirrors positioned just beneath the objective. A 785 nm IR beam was used to maintain focus throughout imaging (Crawford et al., 2013; Friedman and Gelles, 2015). The flow chambers, slide passivation and fiducial markers for stage drift correction have previously been described in (Rosen et al., 2020). After the slide surface was coated with 0.01 mg/mL streptavidin (Thermofisher Scientific 21122), 10 pM of the biotinylated/AF488-labeled DNA template was introduced along with an oxygen scavenging system. The oxygen scavenging system (Crawford et al., 2008), used to minimize photobleaching, contains protocatechuate dioxygenase (Sigma P8279) at a final concentration of 0.9 units/ml and protocatechuate acid (Sigma 03930590) at a final concentration of 5 mM. DNA images were acquired with the 488 nm laser at 1.2 mW. In the experiments with two different DNA templates tethered to the same slide surface, they were sequentially introduced and two DNA images were acquired: one with the fluorescent spots of the first DNA template and the other with fluorescent spots of both DNA templates. After acquiring DNA images, PIC assembly reactions were performed in a flow chamber at room temperature in a mixture in transcription buffer (100 mM HEPES (pH 7.6), 500 mM potassium acetate, 25 mM magnesium acetate, 5 mM EDTA, 1 mg/mL BSA) containing yeast nuclear extract, Gal4-VP16 or Gal4-Gcn4, and other reagents with the following final concentrations: extract (6-8 mg protein/mL), Gal4-VP16 or Gal4-Gcn4 (340 nM, (Joo et al., 2019)), oxygen scavenging system, triplet state quenchers (Dave et al., 2009) (1 mM Trolox (Sigma # 238813), 1 mM propyl gallate (Sigma #02370), and 2 mM 4-nitrobenzyl alcohol (Sigma #N12821)), 20 nM Cy5-TMP (Hoskins et al., 2011), 20 µM acetyl-CoA (Sigma #A2056), *E. coli* genomic DNA (0.02 µg/µL), and an ATP depletion system (20 mM glucose and 2 units hexokinase (Sigma H4502)). In the experiments carried out in the presence of 4 NTPs, 400 µM each ATP, UTP, CTP and GTP were added and the ATP-depletion system was omitted.

Successive images in each channel were captured every 1.4 seconds (0.5 s/frame for each channel, plus switching times) over a time course of 800-1200 seconds. Laser powers were 800 µW and 600 µW for the 532 nm and 633 nm laser, respectively. The cited laser powers were measured at an intermediate point in the excitation beam path before the objective lens. Custom software Glimpse implemented with LabView (National Instruments; Austin, TX) operated microscope, laser shutters, filter wheels, the camera and image acquisition (https://github.com/gelles-brandeis/Glimpse).

#### General image analysis procedure

Image data analysis was performed with custom software implemented with MATLAB (The MathWorks; Natick, MA) as previously described (Friedman and Gelles, 2015) https://github.com/gelles-brandeis/CoSMoS_Analysis. Briefly, DNA locations were identified by automatically selecting fluorescent spots in each DNA image followed by manual removal of overlapping spots. Off DNA locations were also selected in the same image by automatically picking areas that do not contain fluorescent spots. The data were corrected for spatial stage drift that occurred during the experiment. The images from the same field of view created at different channels (red, green and blue channels) were spatially mapped by correcting for spatial displacement between the channels. This correction was done using hundreds of reference fluorescent spot pairs. The reference spots were obtained using surface-immobilized oligonucleotides that were labeled with all three dye molecules (AF488, Cy3, and Cy5). Fluorescence emission from the protein channels (red or green channel) at the pre-defined locations corresponding to template DNA spots was integrated from ∼0.16 µm^2^ areas (3 × 3 pixels) to obtain fluorescence intensity time records at each DNA location (**Fig. 2A, 2D**, **3A**, **4A**, **4B, 5C, S3B-C, S4C,** and **S6D**). Images of protein fluorescence at each DNA location were scored to determine the presence or absence of a fluorescent spot, indicating a “bound” or “unbound” state, respectively. Rastergrams of bound and unbound states were plotted using custom MATLAB scripts (**Fig. 2B, 5B** and **S2A**).

### QUANTIFICATION AND STATISTICAL ANALYSIS

#### Rate constants of PIC factor association with DNA

To analyze the DNA-specific apparent first-order rate constants for factor associations with the DNA template (**Fig. 2C**, **S2D-E, S3D**, and **S4B)**, we performed the analysis as described in (Friedman and Gelles, 2015; Rosen et al., 2020). Briefly, we assumed that not all DNA molecules were capable of factor binding. We termed the fraction of DNA molecules that are capable of factor binding to be the active fraction (*A_f_*). We used the times to first binding at each DNA to minimize the artefacts caused by photobleaching and/or photoblinking. We also accounted for nonspecific binding of factor to the flow chamber surface. The nonspecific association rate constant (*k_ns_*) was first determined by analyzing off DNA locations and we then measured the DNA-specific apparent first-order association rate constant (*k_on_*) by fitting the wait intervals from the DNA locations to a model that included both exponential nonspecific binding (with *k_ns_* held fixed at the previously determined value) and exponential specific binding. Standard errors of fit parameters were determined by bootstrapping. Fit parameters are reported in **Table S4**.

#### DNA-specific frequency of factor association with DNA

##### *Total binding frequencies* (right panel in Fig. 2E (right), 3G, and S5A)

We measured the total binding frequency by dividing the total number of binding events by the sum of absent times at all locations. To determine the DNA-specific total binding frequency, we subtracted the total binding frequency at off-DNA locations from that at DNA locations. The vertical axis intercept of the cumulative distribution of dwell interval frequencies (**Fig. 2G**, **3H**, **4E**, and **S2F**) corresponds to the DNA-specific total binding frequency. Standard error in counting was calculated as the standard deviation of a binomial distribution 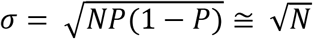 (*if P is close to* 0 *or* 1) where *N* is the total number of observed binding events, *P* is the event probability in time, and σ is the standard deviation.

##### Binding frequencies of GTFs during presence or absence of Pol II (Fig. 3C and 4C)

To examine whether GTF binding preferentially occurs when Pol II is already bound, the DNA-specific binding frequencies of GTFs during Pol II presence or absence were calculated. The Pol II presence time was defined as the sum of the intervals when Pol II, but not the GTF of interest, was present. The binding frequency was calculated as the number of GTF binding events occurring during Pol II presence divided by the Pol II presence time. The binding frequency during Pol II absence was similarly determined by dividing the number of GTF binding events occurring during Pol II absence by the total Pol II absence time. The Pol II absence time was defined as the sum of the intervals when neither Pol II nor the GTF of interest was present. Apparent simultaneous arrival events were not included in this analysis. To determine the DNA-specific binding frequencies, the binding frequencies at off DNA locations were subtracted from those at DNA locations. Standard errors were calculated as the standard deviation of a binomial distribution. We also applied this analysis to determine the DNA-specific binding frequencies of TFIIE during presence or absence of TFIIF (**Fig. 4C**).

#### Cumulative distributions of factor dwell times (Fig. 2G, 3H, 4E, and S2F-G)

To compare the dwell intervals of DNA·factor complexes between the UAS and UAS+promoter DNA templates, we plotted the cumulative DNA-specific binding frequency distributions for the UAS and UAS+promoter DNA. To visually examine whether two cumulative distributions are statistically different, we generated 1000 bootstrap sample sets of the experimental data for the DNA and off DNA locations. We subtracted the off DNA binding frequencies from the DNA binding frequencies and plotted the 90% confidence interval envelope of the bootstrapped samples.

#### Analyzing orders of factor addition during complex assembly (Fig. 3B, 3E, 4F, 4G, 5D, S5D, and S6B)

Cy5- and DY549-labeled factors (i.e., GTF^DHFR-Cy5^ and Rpb1^SNAPf-DY549^) were imaged by alternating laser excitation at 633 nm and 532 nm. The time between two consecutive frames in the same channel was ∼1.4 s (0.5 s/frame for each channel, plus switching times). To analyze the order of binding of Cy5- and DY549-labeled factors to DNA during their complex formation, time intervals in which both factors were present simultaneously for at least part of their bindings to the same DNA were scored as a colocalization. Only de novo colocalizations were included for this analysis. For the selected events, the delay times between arrival times of Cy5- and DY549-labeled factors (e.g. *t*_TFIIF_ –*t*_Pol II_) were determined. Probability density histograms of the delay times were plotted (e.g., **Fig. 3B**, *top*). Standard errors of the bar heights were calculated from the binomial distribution. The events with a time difference between −1.4 s and +1.4 s (± 1 frame) were scored as apparent simultaneous arrivals and were put in the same bin (e.g., **Fig. 3B**, orange bar).

To rule out the possibility that colocalization of two factors on the same DNA molecules occurred by chance, we generated the control scrambled data by performing simulations in which the time record of protein fluorescence from each DNA location was randomly paired with the record of the second protein taken from a different DNA location. We performed 30-50 such simulations, yielding 2000-3000 coincident appearances of two factors. These scrambled control data were then analyzed in the same way as the experimental data.

We classified the complex assembly pathway based on the order of factor binding. Events with a delay time greater than 1.4 s (+1 frame) were scored as a first arrival of the DY549-labeled factor (e.g., **Fig. 3B**, purple bars). Events with a delay time less than -1.4 s (-1 frame) were scored as a first arrival of the Cy5-labeled factor (e.g., **Fig. 3B**, yellow bars).

#### Estimating the fraction of Pol II and TFIIE sequentially arriving at DNA within the experimental time resolution (Fig. S5C)

We fit the part of **Fig. 4F** delay time distribution where Pol II arrival is followed by TFIIE arrival using a maximum likelihood approach that accounts for contributions by both sequential and simultaneous arrival events. Delay times between Pol II and TFIIE arrivals are modeled using a bi-exponential distribution and we account for the fraction of simultaneous arrivals by including a parameter *S* in the equation (1) expression for the likelihood *L*(*a*, *r_1_*, *r_2_*, *S*).

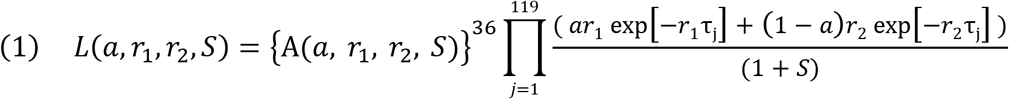

where

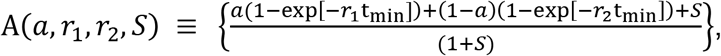

*a* is the relative amplitude, *r_1_* and *r_2_* are two characteristic rates, and *S* is the ratio of simultaneous to sequential arrival events.

In this data set we recorded 119 intervals of duration τ_j_ > *t*_min_, and these data arise from arrival of Pol II followed by a TFIIE arrival with Pol II still present. Each factor of (*ar*_1_ exp[−*r*_1_τ_j_] + (1 − a)*r*_2_ exp[−*r*_2_τ_j_]) / (1 + *S*) in equation (1) is proportional to the probability of observing one such interval of duration τ_j_ separating the sequential arrival of Pol II and TFIIE. Alternating image acquisition for the two proteins means that initial detections of Pol II and TFIIE that are separated by only one frame (separation interval τ<*t*_min_) may be due to either a simultaneous or sequential landing by the Pol II and TFIIE. This data set contained 36 such events, and each *A*(*a*, *r_1_*, *r_2_*, *S*) factor in equation (1) is the probability of observing one such event for which the Pol II/TFIIE landings might have been either simultaneous or sequential. The *a*, *r_1_* and *r_2_* parameters for the bi-exponential distribution describing sequential Pol II/TFIIE landings along with the *S* parameter are determined by optimizing the value of the likelihood function *L*(*a*, *r_1_*, *r_2_*, *S*) in equation (1). The number of sequential arrival events was calculated as 155/(1+*S*) and the number of simultaneous arrival events was calculated as (155 *S*) / (1+*S*), where 155 is the total number of observations in this data set. The standard errors of the *a*, *r_1_*, *r_2_* and *S* fit parameters were determined by bootstrapping.

#### Recruitment rate constant analysis (Fig. 3D)

To determine the recruitment rate constant of TFIIF by pre-bound Pol II, the number of TFIIF bindings occurring during Pol II presence (purple bars in Figure 3B) was divided by the total dwell times of Pol II up to the point, if any, that TFIIF bound. Only the first TFIIF binding at each Pol II was included. The recruitment rate constant determined here may underestimate the true recruitment rate constant of TFIIF by pre-bound Pol II since the sequential binding of Pol II followed by TFIIF within the experimental time resolution were not included. Similar analysis was performed for the scrambled control data.

## Supplemental Information

**Table S1:**
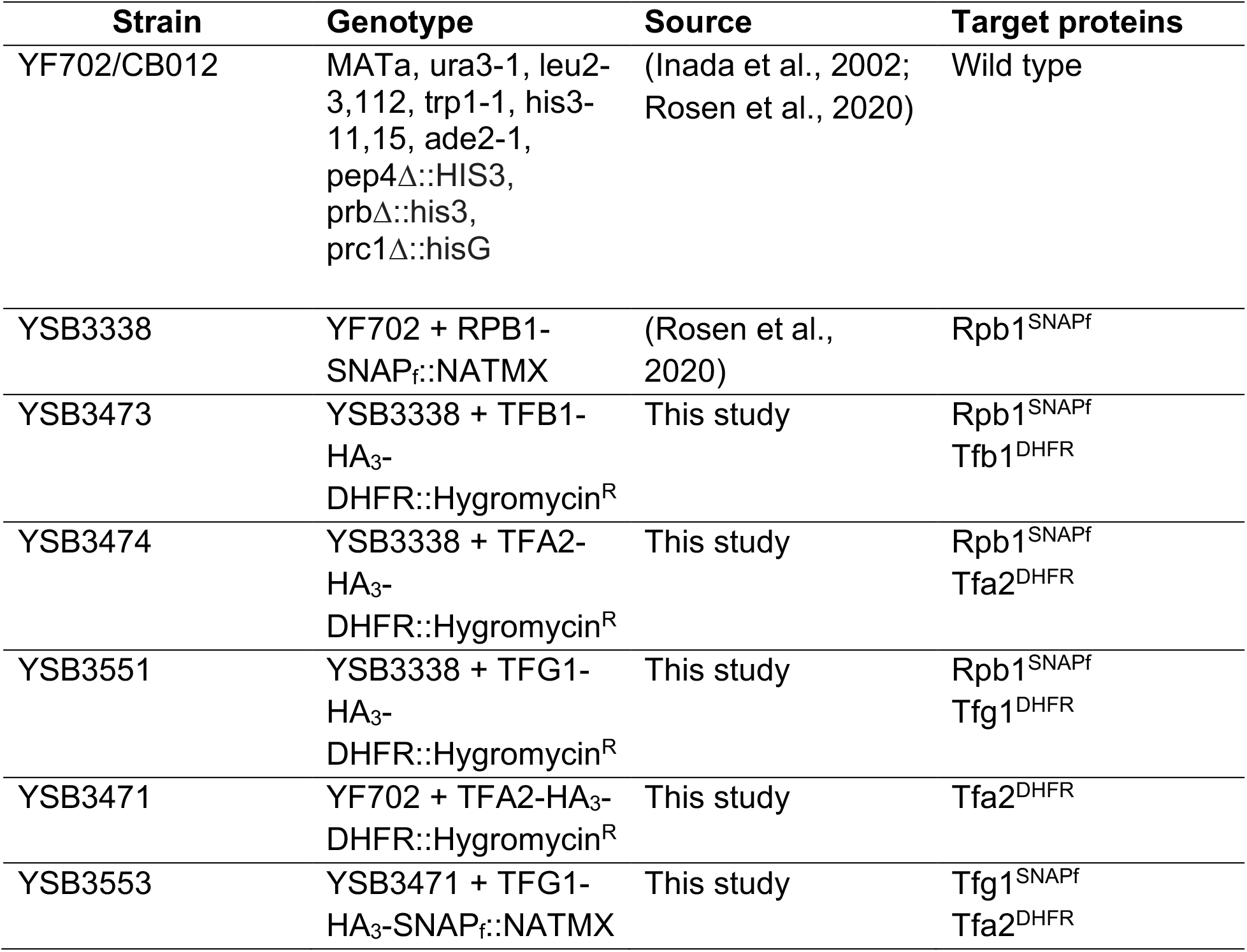
*S. cerevisiae* strains used in this study.

| **Strain** | **Genotype** | **Source** | **Target proteins** |
| --- | --- | --- | --- |
| YF702/CB012 | MATa, ura3-1, leu2-3,112, trp1-1, his3-11,15, ade2-1, pep4Δ::HIS3, prbΔ::his3, prc1Δ::hisG | (Inada et al., 2002; Rosen et al., 2020) | Wild type |
| YSB3338 | YF702 + RPB1-SNAPf::NATMX | (Rosen et al., 2020) | Rpb1<sup>SNAPf</sup> |
| YSB3473 | YSB3338 + TFB1-HA<sub>3</sub>-DHFR::Hygromycin<sup>R</sup> | This study | Rpb1<sup>SNAPf</sup><br>Tfb1<sup>DHFR</sup> |
| YSB3474 | YSB3338 + TFA2-HA<sub>3</sub>-DHFR::Hygromycin<sup>R</sup> | This study | Rpb1<sup>SNAPf</sup><br>Tfa2<sup>DHFR</sup> |
| YSB3551 | YSB3338 + TFG1-HA<sub>3</sub>-DHFR::Hygromycin<sup>R</sup> | This study | Rpb1<sup>SNAPf</sup><br>Tfg1<sup>DHFR</sup> |
| YSB3471 | YF702 + TFA2-HA<sub>3</sub>-DHFR::Hygromycin<sup>R</sup> | This study | Tfa2<sup>DHFR</sup> |
| YSB3553 | YSB3471 + TFG1-HA<sub>3</sub>-SNAPf::NATMX | This study | Tfg1<sup>SNAPf</sup><br>Tfa2<sup>DHFR</sup> |

**Table S2:** Plasmids used in this study.

| **Plasmids** | **Description** | **Source** |
| --- | --- | --- |
| pAAH34 | Fast SNAP gene with yeast nourseothricin resistance gene | (Hoskins et al., 2011) |
| pAAH2 | *E. coli* DHFR gene with yeast hygromycin B resistance gene | (Hoskins et al., 2011) |
| pAAH13 | Fast SNAP gene with yeast hygromycin B resistance gene | (Hoskins et al., 2011) |
| pAAH6 | SNAP gene with yeast nourseothricin resistance gene | (Hoskins et al., 2011) |
| pFA6a-HA-KIURA3 | Triple HA tag with URA3 gene | (Sung et al., 2008) |
| pBS-SKII-3XHA-eDHFR-Hygromycin (YV317) | Triple HA tag-*E. coli* DHFR gene with yeast hygromycin B resistance gene | This study |
| pBS-SKII-3XHA-fSNAP-NAT (YV309) | Triple HA tag-fast SNAP gene with yeast nourseothricin resistance gene | This study |
| pUC18-G5CYC1 G- (SB649) | Five copies of GAL4 binding site upstream of *CYC1* promoter driving a G-less cassette | (Johnson et al., 2009) |
| pUC18-G5-HIS4 G- (SB1964) | Five copies of GAL4 binding site upstream of *HIS4* promoter driving a G-less cassette | This study |
| pBlusScript | Plasmid backbone used to generate YV317 and YV309 | Stratagene #212205 |

**Table S3:**
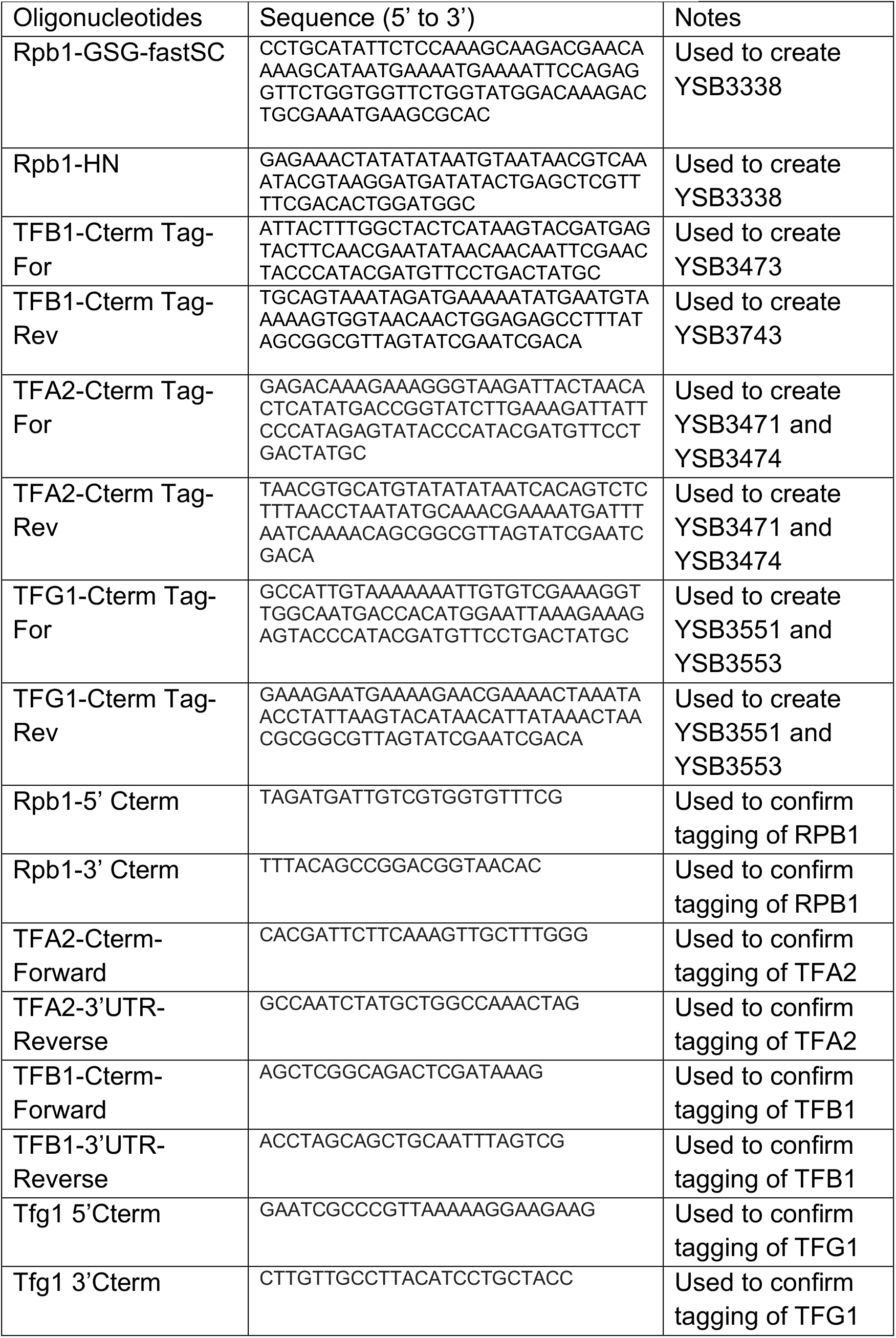
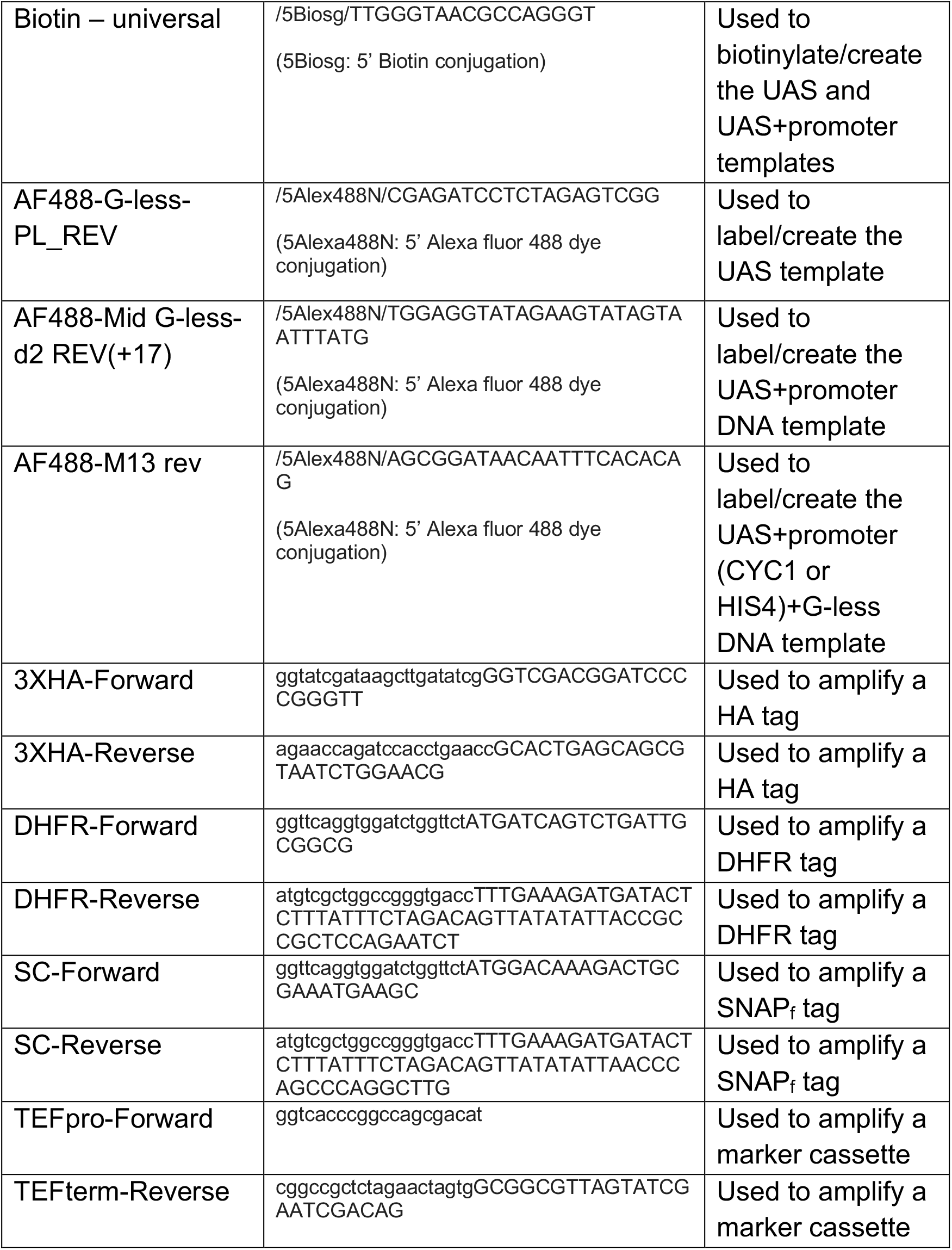
Oligonucleotides used in this study.

**Table S4:** Kinetics for association of labeled factors with DNA.

| **Labeled factor (Extract)** | **Fits** | **DNA template** | **$A_f$** | **$k_{on} (\times 10^{-3} s^{-1})$** | **$k_{ns} (\times 10^{-3} s^{-1})$** |
| --- | --- | --- | --- | --- | --- |
| Rpb1<sup>SNAPf-Dy549</sup><br>(YSB3474) | Fig. 2C | UAS + promoter | $0.73 \pm 0.06$ | $3.8 \pm 1.0$<br>($N = 146$) | $0.22 \pm 0.02$<br>($N = 150$) |
| Rpb1<sup>SNAPf-Dy549</sup><br>(YSB3474) | Fig. 2C | UAS | $0.78 \pm 0.04$ | $4.4 \pm 0.5$<br>($N = 165$) | $0.22 \pm 0.02$<br>($N = 150$) |
| Rpb1<sup>SNAPf-Dy549</sup><br>(YSB3551) | - | UAS + promoter | $0.61 \pm 0.05$ | $4.9 \pm 1.2$<br>($N = 92$) | $0.10 \pm 0.01$<br>($N = 78$) |
| Rpb1<sup>SNAPf-Dy549</sup><br>(YSB3551) | - | UAS | $0.54 \pm 0.03$ | $6.8 \pm 1.1$<br>($N = 218$) | $0.10 \pm 0.01$<br>($N = 78$) |
| Rpb1<sup>SNAPf-Dy549</sup><br>(YSB3473) | - | UAS + promoter | $0.90 \pm 0.02$ | $5.5 \pm 0.8$<br>($N = 178$) | $0.07 \pm 0.01$<br>($N = 66$) |
| Rpb1<sup>SNAPf-Dy549</sup><br>(YSB3473) | - | UAS | $0.88 \pm 0.02$ | $5.9 \pm 0.5$<br>($N = 404$) | $0.07 \pm 0.01$<br>($N = 66$) |
| Rpb1<sup>SNAPf-Dy549</sup><br>(YSB3473) | Fig. S2D<br>Fig. S2E<br>(left) | UAS+HIS4-G-less | $0.70 \pm 0.05$ | $8.8 \pm 1.4$<br>($N = 156$) | $1.3 \pm 0.1$<br>($N = 441$) |
| Rpb1<sup>SNAPf-Dy549</sup><br>(YSB3473) | Fig. S2D<br>Fig. S2E<br>(left) | UAS | $0.69 \pm 0.05$ | $11.9 \pm 2.7$<br>($N = 229$) | $1.3 \pm 0.1$<br>($N = 441$) |
| Rpb1<sup>SNAPf-Dy549</sup><br>(YSB3473) | Fig. S2E<br>(right) | UAS+HIS4-G-less | $0.76 \pm 0.06$ | $7.3 \pm 1.4$<br>($N = 157$) | $1.3 \pm 0.1$<br>($N = 414$) |
| Rpb1<sup>SNAPf-Dy549</sup><br>(YSB3473) | Fig. S2E<br>(right) | UAS+CYC1-G-less | $0.76 \pm 0.03$ | $7.8 \pm 1.0$<br>($N = 327$) | $1.3 \pm 0.1$<br>($N = 414$) |
| Tfg1<sup>DHFR-Cy5</sup><br>(YSB3551) | Fig. 3F<br>Fig. S3D | UAS + promoter | $0.54 \pm 0.09$ | $3.5 \pm 1.9$<br>($N = 110$) | $0.56 \pm 0.03$<br>($N = 331$) |
| Tfg1<sup>DHFR-Cy5</sup><br>(YSB3551) | Fig. 3F<br>Fig. S3D | UAS | $0.45 \pm 0.04$ | $4.7 \pm 0.9$<br>($N = 258$) | $0.56 \pm 0.03$<br>($N = 331$) |
| Tfa2<sup>DHFR-Cy5</sup><br>(YSB3474) | Fig. 4D<br>Fig. S4B | UAS + promoter | $0.54 \pm 0.12$ | $2.3 \pm 2.1$<br>($N = 147$) | $0.75 \pm 0.04$<br>($N = 362$) |
| Tfa2<sup>DHFR-Cy5</sup><br>(YSB3474) | Fig. 4D<br>Fig. S4B | UAS | 0.51 ±<br>0.09 | 2.5 ± 0.7<br>(*N* = 157) | 0.75 ± 0.04<br>(*N* = 362) |

\* $k_{on}$ and $k_{ns}$ are the apparent first-order rate constants for binding to DNA and for non-specific binding to the chamber surface.
$A_f$ is the active fraction of DNA molecules capable of factor binding.
$N$ is the number of DNA or off DNA sites with at least one binding.
1000 bootstrap iterations were performed to determine standard errors of fit parameters.

**Table S5:**
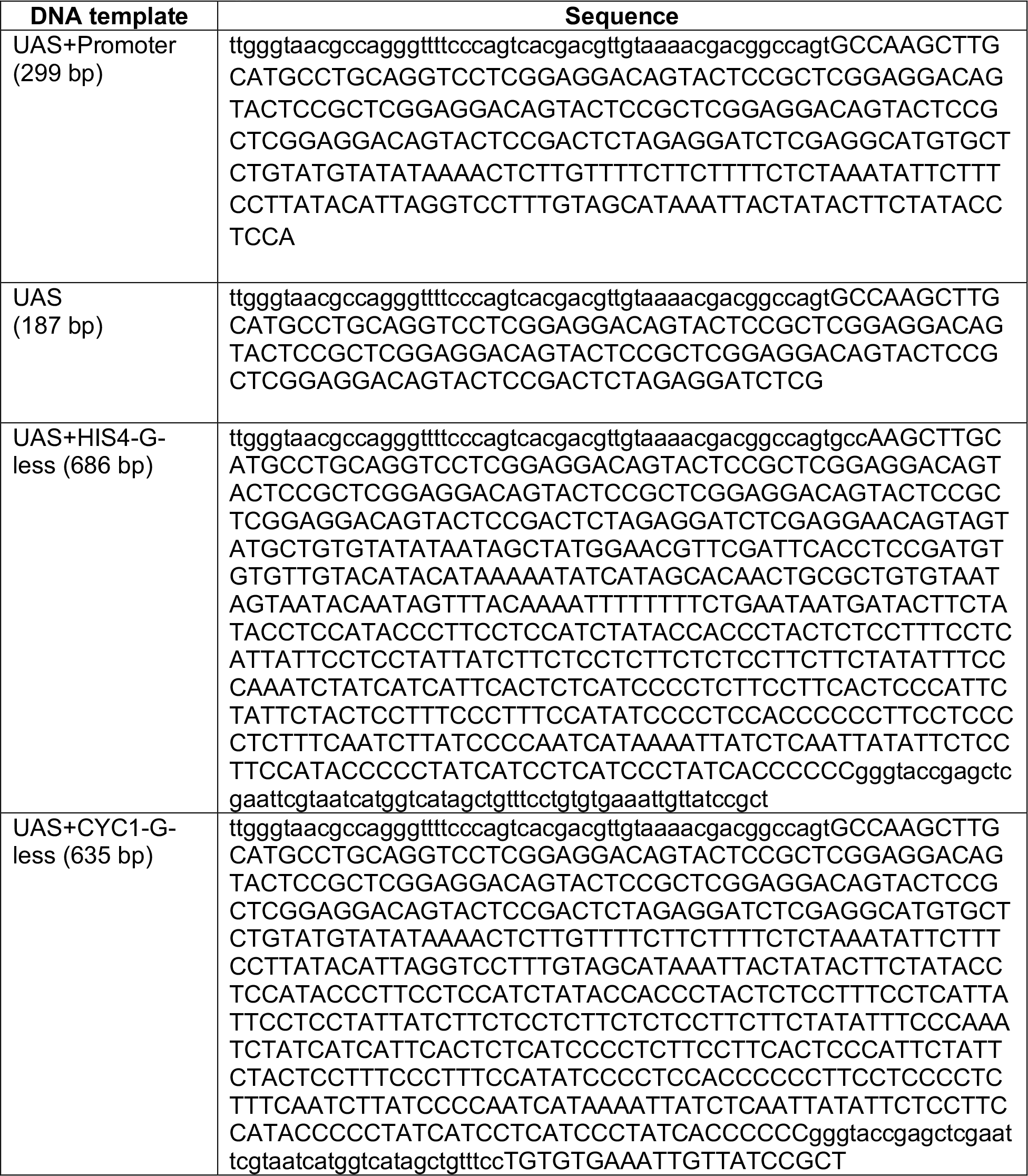
DNA templates.

**Figure S1, related to Figure 1.**
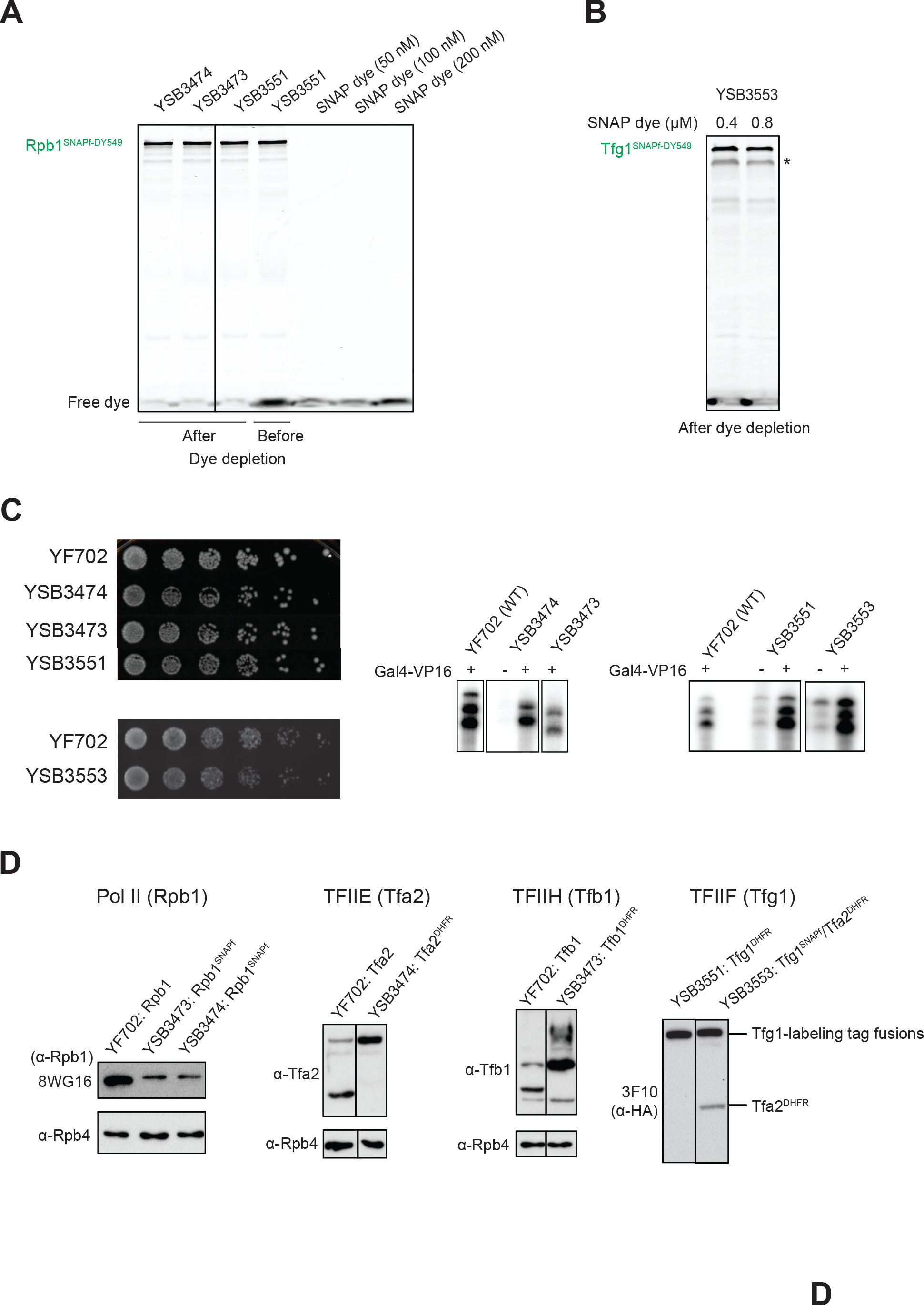
Single molecule imaging of Pol II and GTF binding to the DNA template. **(A-B)** SDS (sodium dodecyl sulfate)-PAGE (polyacrylamide gel electrophoresis) gel followed by fluorescence imaging for SNAP-surface 549 dye. **(A)** Three different nuclear extracts (lanes 1-3) containing Rpb1^SNAPf-DY549^ prepared as previously described(Rosen et al., 2020) (also see Method details). Rpb1-SNAP_f_ was labeled with SNAP-Surface 549 dye with high specificity in all three nuclear extracts. The concentrations of residual dye in each nuclear extract after dye depletion are less than 50 nM. (SNAP dye: SNAP-Surface 549, YSB3473: Rpb1^SNAPf^/Tfb1^DHFR^ strain, YSB3474: Rpb1^SNAPf^/Tfa2^DHFR^, and YSB3551: Rpb1^SNAPf^/Tfg1^DHFR^ strain; see **Table S1**). Lane 4 shows the extract from YSB3551 before dye depletion. **(B)** Tfg1^SNAPf-DY549^/Tfa2^DHFR^ nuclear extract prepared with two different concentrations of SNAP-Surface 549 dye and depleted of unreacted dye after extract labeling. A concentration of 0.4 µM SNAP-Surface 549 was sufficient for maximal labeling of Tfg1-SNAP_f_ in nuclear extract. (asterisk: presumed to be a cleaved fragment of Tfg1^SNAPf-DY549^ generated during the nuclear extract preparation; YSB3553: Tfg1^SNAPf^/Tfa2^DHFR^ strain; see **Table S1**) **(C)** (*left*) Yeast spot assay for visualizing cell growth. 4-fold serial dilutions were grown on YPD medium at 30°C. The fusion strains exhibited no growth defect in comparison with YF702 (untagged parent strain; see **Table S1**). (*right*) Bulk *in vitro* transcription assay for measuring RNA transcripts produced from plasmid templates through transcription in nuclear extracts from YF702 and the fusion strains. Nuclear extracts from the fusion strains were able to produce RNA transcripts in an activator-dependent manner to a similar level observed in nuclear extract from YF702. **(D)** Western blot of whole cell lysates showing that the labeling tag fusions did not perturb protein expression levels. Rpb4 was used as a loading control. Rpb1 was detected by monoclonal antibody 8WG16(Thompson et al., 1990), Rpb4 by a polyclonal anti-Rpb4 antibody (Biolegend #665106), Tfa2 by a polyclonal anti-Tfa2 antibody (Kuldell and Buratowski, 1997), and Tfb1 by a polyclonal anti-Tfb1 antibody(Matsui et al., 1995). Due to a lack of an anti-Tfg1 antibody, Tfg1^SNAPf^ or Tfg1^DHFR^ was assessed with the anti-HA-peroxidase antibody 3F10 (Roche, #12013819001). **(A-D)** Irrelevant lanes are eliminated and the deletion positions are indicated with a line.

**Figure S2, related to Figure 2.**
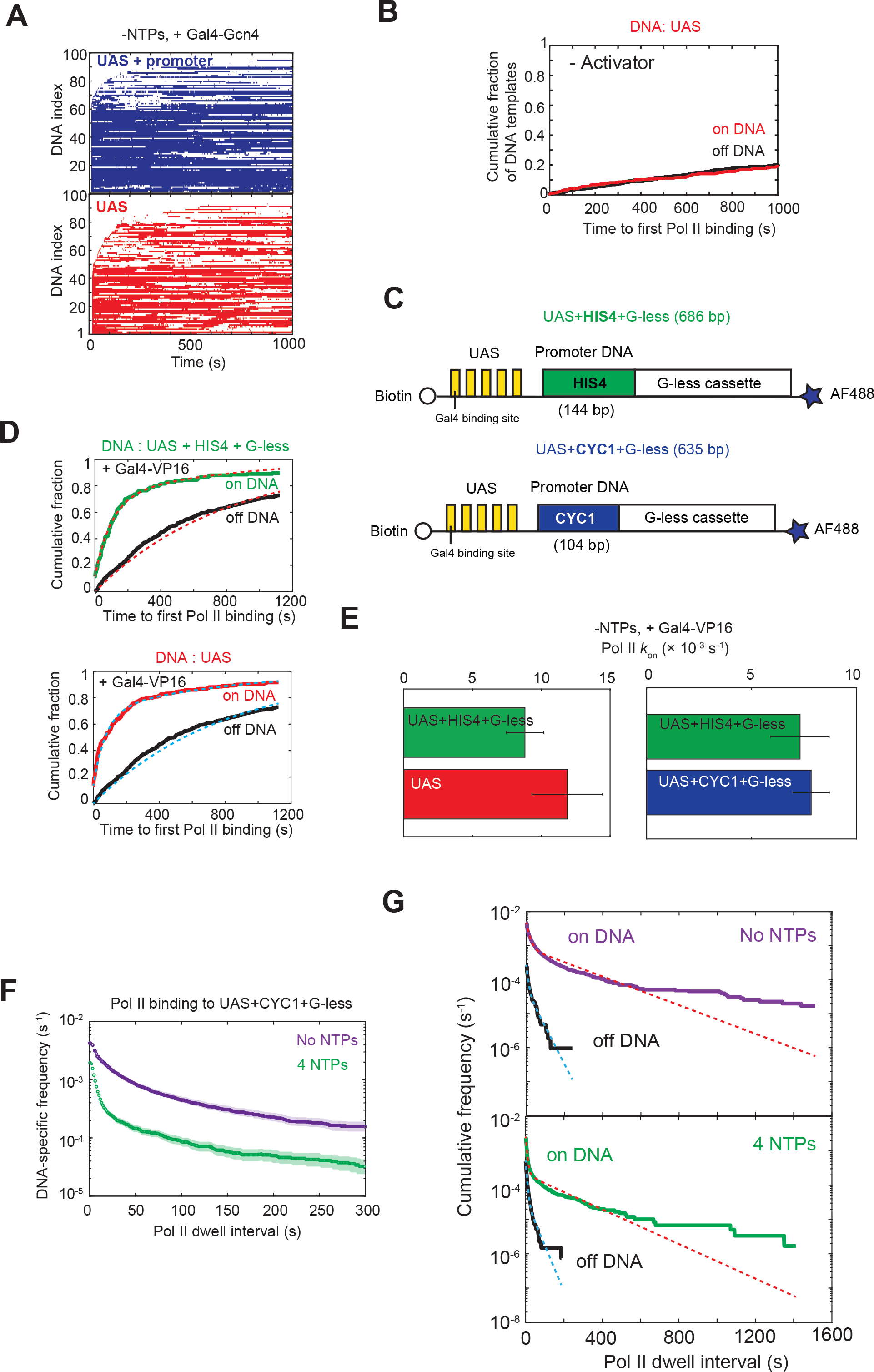
Pol II association kinetics are independent of the core promoter. **(A)** Rastergrams of Pol II binding to 100 randomly selected DNA locations, plotted as in **Fig. 2B**. **(B)** Cumulative distributions over time for the fraction of UAS alone DNA molecules (red) and the fraction of off DNA sites (black) bound at least once by Pol II in the absence of activator. Off DNA locations are control locations that do not contain DNA fluorescence spots. No Pol II binding to the UAS above background was observed in the absence of activator. **(C)** Schematic of the DNA templates used for examining the effects of the promoter element or NTP addition on Pol II association and dissociation kinetics. The DNA template contains five Gal4-binding sites (yellow) and the *CYC1* (blue) or *HIS4* (green) core promoter sequence followed by a G-less cassette of ∼300 bp. The 5’ end is biotinylated (empty circle) and the 3’ end is labeled with Alexa Fluor 488 dye (AF488, blue star). **(D)** Cumulative fraction of DNA molecules bound at least once by Pol II as a function of time (*top*: UAS+HIS4-G-less (green), *bottom*: UAS (red)). Binding to off DNA sites (black) is shown as a control for background. Data were fit to a single exponential model (red or cyan dashed lines). Fit parameters and number of observations are given in panel E (left) and **Table S4**. **(E)** Apparent first-order rate constants (±S.E.) of Pol II initial association with the UAS+HIS4-G-less (green), UAS (red), or UAS+CYC1-G-less (blue) calculated from fitting to a single exponential model. (Fit parameters in **Table S4**). Left and right graphs are from separate experiments. **(F)** Cumulative distributions of Pol II dwell intervals on the UAS+CYC1+G-less DNA templates with 90% confidence intervals in the absence of NTPs (purple) or in the presence of 4 NTPs (green) when Gal4-VP16 was present. As in **Fig. 2D**, each dwell consists of a continuous time interval in which one or more labeled Pol II molecules were present on the DNA. Frequency values on the vertical axis are after subtraction of off DNA background. **(G)** Same Pol II dwell interval data in (F) were used, but frequency values on the vertical axis were before subtraction of off DNA background (purple, no NTPs; green, 4 NTPs) to show on and off DNA fit curves. Cumulative distributions of Pol II dwell intervals on off DNA locations were also plotted (black curves). (*top*) Pol II dwell interval data in the absence of NTPs were fit to biexponential distributions for both the DNA sites (red; *r*_1_ = (0.7 ± 0.1) × (10^-1^) s^-1^, *r*_2_ = (4.8 ± 0.7) × (10^-3^) s^-1^, and *a* = 0.81 ± 0.02) and the off DNA sites (cyan; *r*_1_ = (1.3 ± 0.1) × (10^-1^) s^-1^, *r*_2_ = (2.5 ± 0.6) × (10^-2^) s^-1^, and *a* = 0.81 ± 0.06). (*bottom*) Pol II dwell interval data in the presence of NTPs were fit using biexponential distributions for both the DNA sites (red; *r*_1_ = (1.6 ± 0.1) × (10^-1^) s^-1^, *r*_2_ = (5.8 ± 1.1) × (10^-3^) s^-1^, and *a* = 0.90 ± 0.01) and the off DNA sites (cyan; *r*_1_ = (1.8 ± 0.1) × (10^-1^) s^-1^, *r*_2_ = (3.1 ± 1.1) × (10^-2^) s^-1^, and *a* = 0.93 ± 0.04). *a* is the relative amplitude and *r*_1_ and *r*_2_ are two characteristic departure rates (Friedman and Gelles, 2015).

**Figure S3, related to Figure 3.**
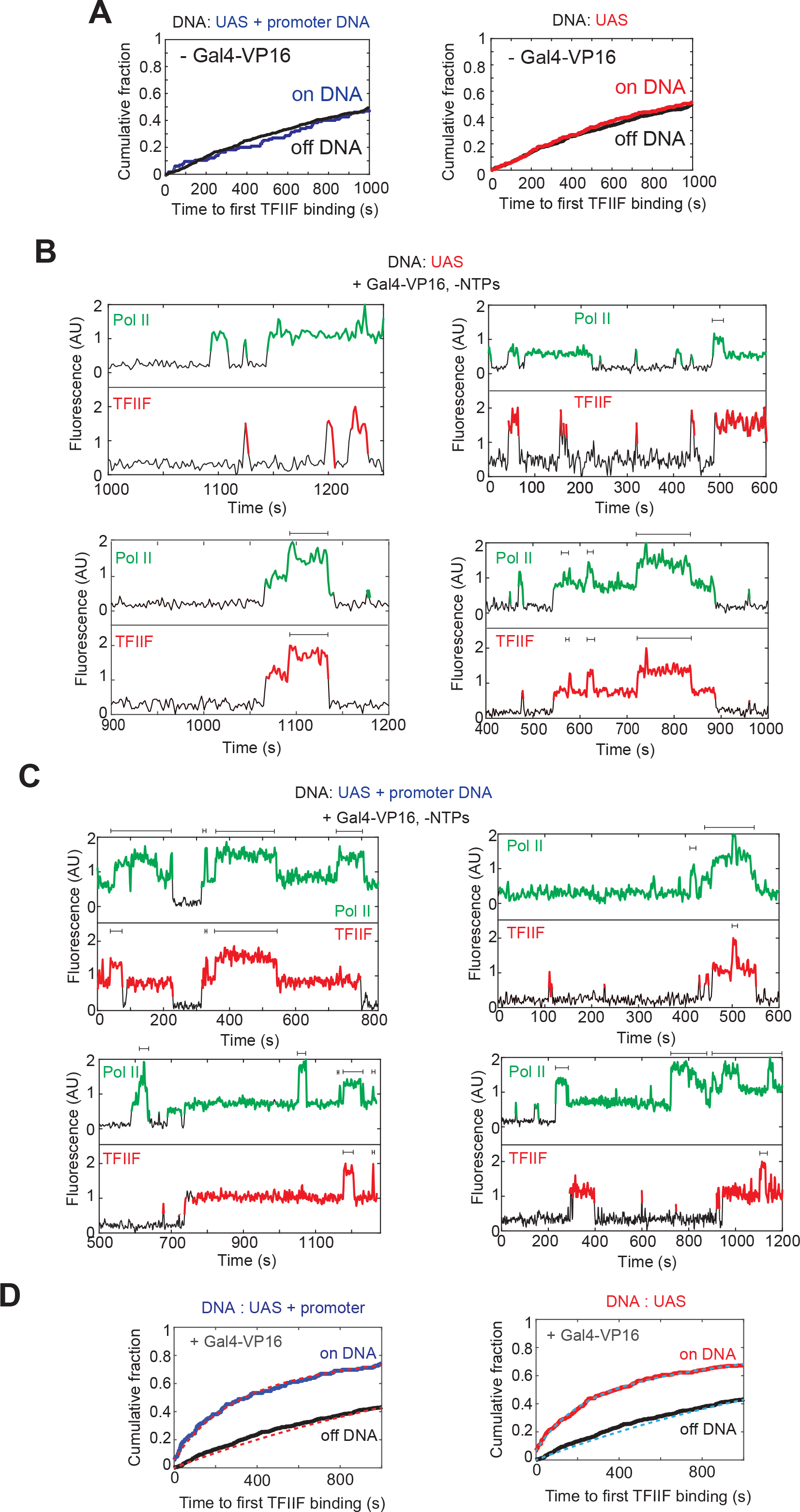
Dynamics of Pol II and TFIIF during activator-dependent PIC assembly. **(A)** Cumulative distributions over time for the fraction of UAS+promoter DNA molecules (*left*, blue) or UAS (*right*, red) and the fraction of off DNA sites (black) bound at least once by TFIIF in the absence of Gal4-VP16. No background above TFIIF binding to the UAS+promoter or UAS was observed in the absence of Gal4-VP16. **(B)** Time records of Pol II fluorescence and TFIIF fluorescence at the locations of four different UAS alone DNA molecules in the presence of Gal4-VP16 and the absence of NTPs. Colored intervals are times at which Pol II (green) or TFIIF (red) was colocalized to the DNA molecules. Brackets on the records indicate times at which multiple Pol II or multiple TFIIF molecules simultaneously bound to the same DNA molecule. The simultaneous occupancy of multiple TFIIF molecules is correlated with that of multiple Pol II molecules. **(C)** Time records of Pol II fluorescence and TFIIF fluorescence at the locations of four different UAS+promoter DNA molecules in the presence of Gal4-VP16 and the absence of NTPs, plotted as in panel B. Note that the presence of multiple TFIIF molecules appears correlated with that of multiple Pol II molecules. **(D)** Same data as in **Fig. 3F**, along with fits (left: red dashed lines and right: cyan dashed lines) to a single exponential binding model. Fit parameters are given in **Table S4**.

**Figure S4, related to Figure 4.**
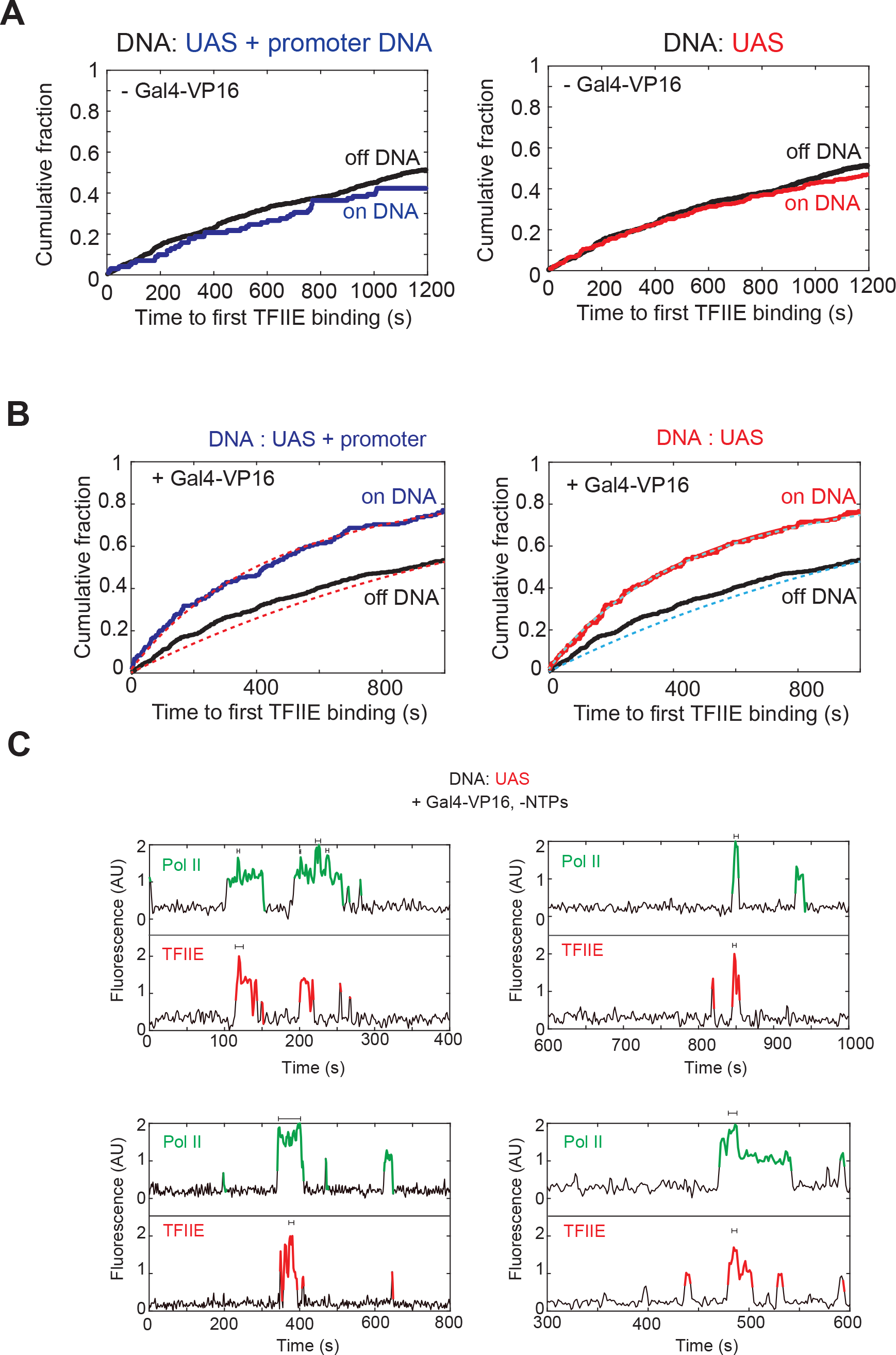
Dynamics of TFIIE relative to Pol II and TFIIF during activator-dependent PIC assembly. **(A)** Cumulative distributions over time for the fraction of UAS+promoter DNA molecules (*left*, blue) or UAS (*right*, red) and the fraction of off DNA sites (black) bound at least once by TFIIE in the absence of Gal4-VP16. **(B)** Same data as in **Fig. 4D**, along with fits (left: red dashed lines and right: cyan dashed lines) to a single exponential binding model. Fit parameters are given in **Table S4**. **(C)** Time records of Pol II fluorescence and TFIIE fluorescence at the locations of four different UAS alone DNA molecules in the presence of Gal4-VP16 and the absence of NTPs. Colored intervals are times at which Pol II (green) or TFIIE (red) was colocalized to the DNA molecules. Brackets on the TFIIE records indicate times at which multiple TFIIE molecules simultaneously bound to the same DNA molecule

**Figure S5, related to Figure 4.**
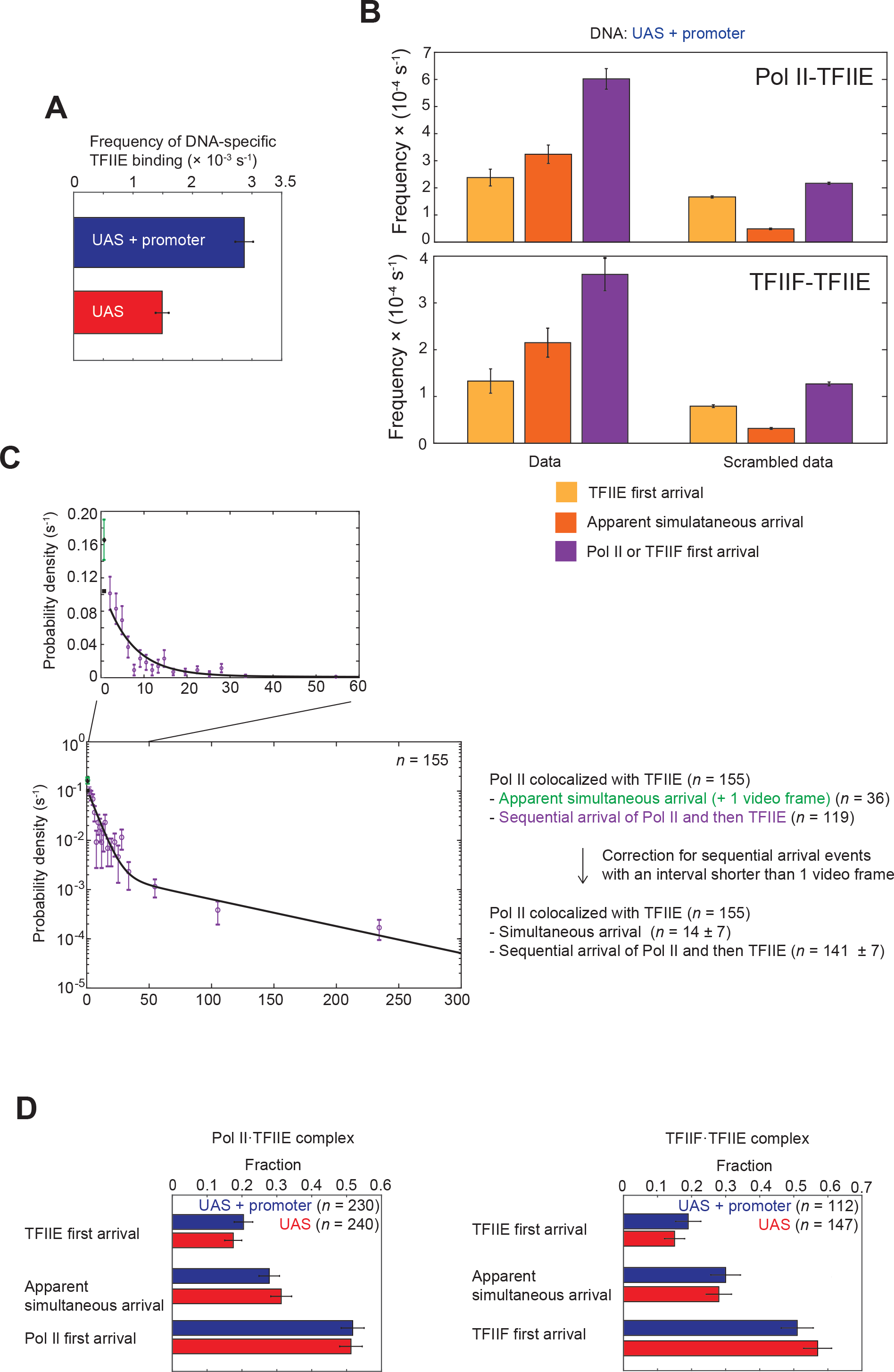
Dynamics of TFIIE relative to Pol II and TFIIF during activator-dependent PIC assembly. **(A)** DNA-specific binding frequencies (±S.E.) of all TFIIE bindings to the UAS+promoter DNA (blue) and the UAS (red). **(B)** Frequencies (±S.E.) of each pathway for the Pol II·TFIIE complex (*top*) and the TFIIF·TFIIE complex (*bottom*). The first three bars are from the data and the last three bars are from the scrambled data. (Yellow bars: TFIIE first arrival events, orange bars: apparent simultaneous arrival events and purple bars: Pol II or TFIIF first arrival events). **(C)** Probability density (±S.E.) distribution of delay times between TFIIE and Pol II arrivals for colocalizations where Pol II was detected before TFIIE (Pol II first arrival, *n* = 155). A subset of the Pol II first arrival events was scored as being due to either simultaneous or rapid sequential arrival of Pol II and TFIIE (green point, *n* = 36), as TFIIIE was present within the first red channel frame after Pol II detection in the green channel. The rest of the Pol II first arrival events were scored as unambiguously being due to the sequential arrival of Pol II and then TFIIE (purple points, *n* = 119). The distribution for the entire data set used a bi-exponential function to model the contribution from sequential arrivals and an offset parameter *S* to account for simultaneous arrivals (Quantification and statistical analysis equation (1)). This treatment yielded fit parameters *r*_1_= 0.15 ± 0.06 s^-1^, *r*_2_ = 0.013 ± 0.005 s^-1^, *a* = 0.80 ± 0.07 and *S* = 0.097 ± 0.048. Black line shows the distribution calculated from the model using these fit parameters from time *t*_min_ to infinity. Square point (first bin) represents a probability density of just sequential arrival events within the experimental time resolution calculated using the fit parameters. Diamond point (first bin) represents a probability density of both sequential and simultaneous events with the experimental time resolution calculated using the fit parameters. After correcting for the sequential bindings predicted to occur within an interval less than the imaging time resolution, 14 ± 7 events are likely to be a simultaneous arrival of Pol II and TFIIE, which accounts for 9 ± 5% of the total events (155 events). **(D) (***left*) Fractions (± S.E.) of the pathways for the formation of Pol II·TFIIE complex on the UAS+promoter DNA (blue) or on the UAS (red). (*right*) Fractions (± S.E.) of the pathways for the formation of TFIIF·TFIIE complex on the UAS+promoter DNA (blue) or on the UAS (red). *n* specifies the total number of the observations in each condition.

**Figure S6, related to Figure 5.**
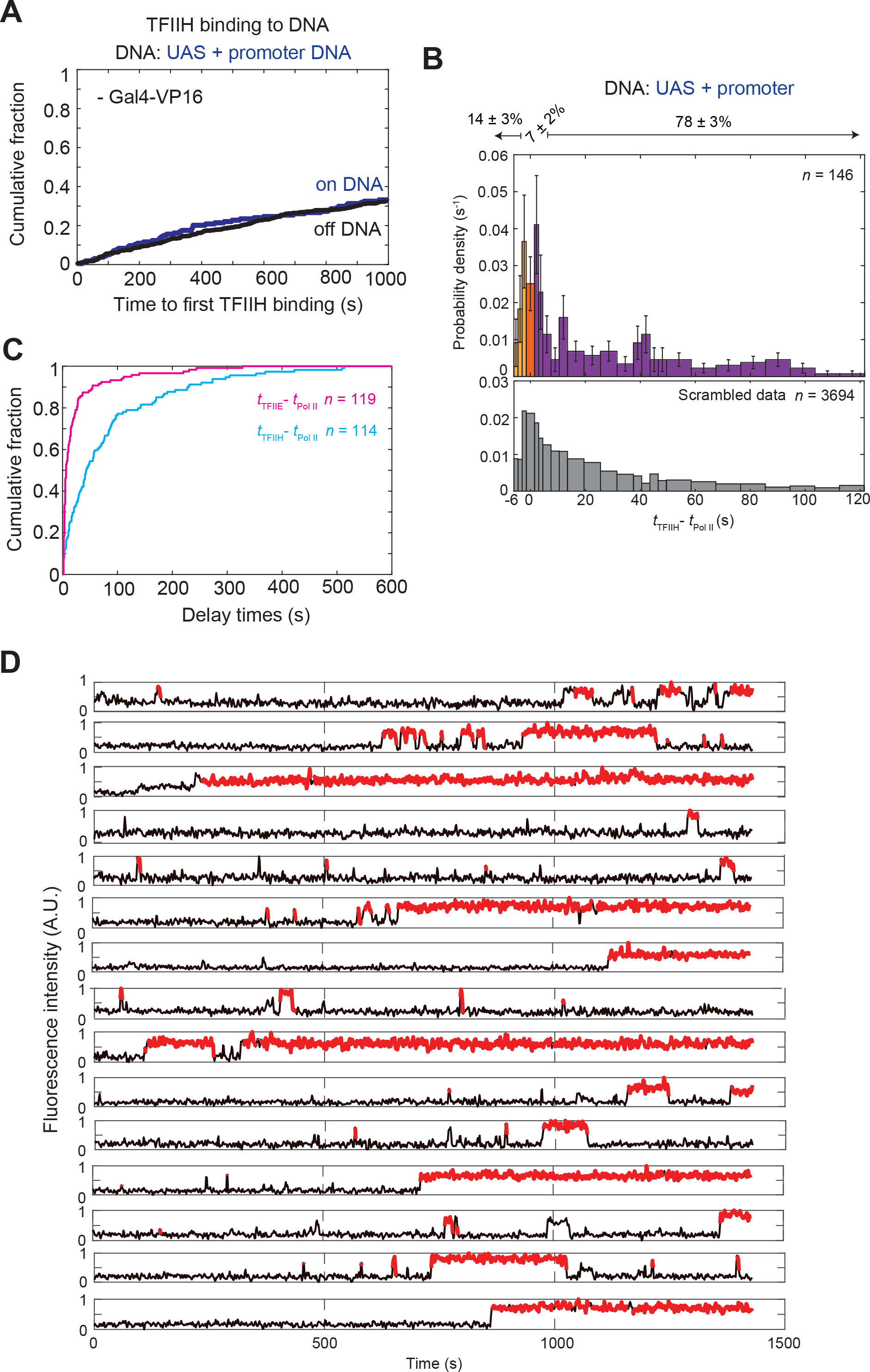
Dynamics of TFIIH relative to Pol II during activator-dependent PIC assembly. **(A)** Cumulative distributions versus time for the fraction of UAS+promoter molecules and the fraction of off DNA sites (black) bound at least once by TFIIH in the absence of Gal4-VP16. No specific TFIIH binding to DNA was observed in the absence of Gal4-VP16. **(B)** Probability density (±S.E.) histogram of the differences between arrival times of TFIIH and Pol II (*t*_TFIIH_ − *t*_Pol II_) at unoccupied UAS+promoter DNA molecules plotted as in **Fig. 3B.** The range shown contains 79% of the measured *t*_TFIIH_ − *t*_Pol II_ values. **(C)** Cumulative distributions of delay times between Pol II and TFIIE arrivals (magenta) or between Pol II and TFIIH arrivals (cyan) leading to colocalization. **(D)** Time records of TFIIH fluorescence at the locations of 15 different UAS+promoter DNA molecules in the presence of Gal4-VP16 and the absence of NTPs. Red-colored intervals are times at TFIIH was colocalized to the DNA molecules.

